# DCLK1-Mediated Regulation of Invadopodia Dynamics and Matrix Metalloproteinase Trafficking Drives Invasive Progression in Head and Neck Squamous Cell Carcinoma

**DOI:** 10.1101/2024.04.06.588339

**Authors:** Levi Arnold, Marion Yap, Laura Jackson, Michael Barry, Thuc Ly, Austin Morrison, Juan P. Gomez, Michael P. Washburn, David Standing, Nanda Kumar Yellapu, Linheng Li, Shahid Umar, Shrikant Anant, Sufi Mary Thomas

## Abstract

Head and neck squamous cell carcinoma (HNSCC) is a major health concern due to its high mortality from poor treatment responses and locoregional tumor invasion into life sustaining structures in the head and neck. A deeper comprehension of HNSCC invasion mechanisms holds the potential to inform targeted therapies that may enhance patient survival. We previously reported that doublecortin like kinase 1 (DCLK1) regulates invasion of HNSCC cells. Here, we tested the hypothesis that DCLK1 regulates proteins within invadopodia to facilitate HNSCC invasion. Invadopodia are specialized subcellular protrusions secreting matrix metalloproteinases that degrade the extracellular matrix (ECM). Through a comprehensive proteome analysis comparing DCLK1 control and shDCLK1 conditions, our findings reveal that DCLK1 plays a pivotal role in regulating proteins that orchestrate cytoskeletal and ECM remodeling, contributing to cell invasion. Further, we demonstrate in TCGA datasets that DCLK1 levels correlate with increasing histological grade and lymph node metastasis. We identified higher expression of DCLK1 in the leading edge of HNSCC tissue. Knockdown of DCLK1 in HNSCC reduced the number of invadopodia, cell adhesion and colony formation. Using super resolution microscopy, we demonstrate localization of DCLK1 in invadopodia and colocalization with mature invadopodia markers TKS4, TKS5, cortactin and MT1-MMP. We carried out phosphoproteomics and validated using immunofluorescence and proximity ligation assays, the interaction between DCLK1 and motor protein KIF16B. Pharmacological inhibition or knockdown of DCLK1 reduced interaction with KIF16B, secretion of MMPs, and cell invasion. This research unveils a novel function of DCLK1 within invadopodia to regulate the trafficking of matrix degrading cargo. The work highlights the impact of targeting DCLK1 to inhibit locoregional invasion, a life-threatening attribute of HNSCC.

## Introduction

Head and neck squamous cell carcinoma (HNSCC) pose a significant global health challenge due to its high incidence rates and adverse effects on patient survival [1]. A key feature of this malignancy is its propensity for locoregional invasion, which greatly contributes to patient mortality and treatment-related complications [2]. The tumor cells’ ability to breach local extracellular matrix (ECM) barriers, invading critical structures such as the carotid artery or metastasizing to distant sites contributes to mortality [3]. Understanding these early metastatic events is crucial for shedding light on HNSCC pathogenesis and identifying potential therapeutic targets in a largely underexplored aspect of HNSCC research.

Invadopodia, actin-rich, finger-like cellular projections, play a pivotal role in cancer cell invasion by puncturing the ECM through the secretion of MMPs [4, 5]. Invadopodia formation involves three stages: initiation, assembly, and maturation [6]. Initiation, stimulated by growth factors or other signals, involves the rapid recruitment of key molecular components, that polymerizes actin filaments near the plasma membrane [7]. In the assembly phase, RGD-binding integrins, integrin-linked kinase, vinculin and paxillin create an adhesion ring around the invadopodia [8]. During the maturation phase, cofilin-driven actin polymerization and Arp2/3-dependent actin nucleation facilitate the development of fully functional invadopodia [9]. Importantly, microtubules elongate into invadopodia, providing structural rigidity and serving as a tract for trafficking vesicles containing MMPs [10]. Additionally, emerging research highlights the localization of adaptor protein, tyrosine kinase substrate with four Src homology 3 domains 4 (Tks4) and recruitment of MT1-MMP to the distal end of the invadopodia [11, 12]. Adaptor protein Tks5, acts as a tether for Rab40b that binds to vesicles containing MMPs and transports they to the elongating invadopodia [13]. The development of invadopodia and molecular mechanisms regulating its phenotype are not fully characterized.

Doublecortin like kinase 1 (DCLK1), originally characterized as a microtubule-binding protein, has predominantly been associated with cancer stem cell properties, particularly in colorectal cancer [14]. Its relevance in cancer research stems from observations, including recent findings from our research group, which highlight that inhibiting or genetically abrogating DCLK1 reduces cancer cells’ ability to degrade and invade through an ECM substrate [15]. Notably, DCLK1’s history extends beyond cancer studies and finds its origins in neurobiology. Early research identified DCLK1’s presence at the growth cones of axons, where it played a pivotal role in neural elongation [16]. Studies by Lipka et al. further elaborated on DCLK1’s function in the transportation of dense core vesicles, notably in collaboration with the kinesin superfamily, particularly members of the Kinesin 3 family [17].

In our current research, we aimed to unravel the intricate relationship between DCLK1 and invasion, employing proteomic analysis. Our investigations unveiled that DCLK1 exerts significant influence over cell-ECM interactions, cell movement, and cell invasion. Furthermore, our findings corroborate earlier work [17] indicating DCLK1’s interaction with members of the Kinesin 3 family, with KIF16B exhibiting a significant proximity to DCLK1. Crucially, our research also demonstrates that the abrogation of DCLK1 results in a reduction in the secretion of ECM degradative cargo by HNSCC cells. These findings collectively underscore the multifaceted role of DCLK1 in cancer invasion, highlighting its significance in advancing our understanding of cancer biology.

## Results

In the context of HNSCC, the impact of DCLK1 on tumor invasion remains a subject that lacks comprehensive understanding. To address this knowledge gap, we undertook a targeted approach by generating shRNA-mediated knockdown of DCLK1 in FaDu cells (Fig. 1 A.). The proteomic and phosphoproteomic signatures (Sup. Table 1) displayed distinct clustering patterns, highlighting substantial differences in the molecular profiles regulated by DCLK1 (Sup. Fig. 1 A and B). These genetically modified cells were subsequently subjected to tandem mass tag spectrometry. Between the shControl cells and the DCLK1 knockdowns, we identified a substantial alteration in protein expression. While 376 proteins were downregulated in the DCLK1 knockdown cells when compared to the controls, 359 proteins showed upregulation (Fig. 1 B). Network ontological analysis of DCLK1-enriched proteins revealed specific interactions related to cell-ECM interaction (Fig. 1 C). Hierarchical analysis further identified terms associated with ECM interactions, cytoskeleton reorganization, and cell adhesion (Fig. 1 D and E). To understand the functional implications of the identified proteomic and phosphoproteomic alterations, we conducted further analysis using the Metascape tool [18]. The results unveiled a diverse array of roles for DCLK1 in various cellular processes. Notably, there was an enrichment of proteins involved in membrane interactions, particularly in the context of trafficking, suggesting DCLK1’s crucial involvement in regulating the transport of cellular components across membranes (Sup. Fig. 2 A, Sup. Table 2). The data also showed that DCLK1 may be involved in cytoskeletal rearrangement, as evidenced by increased expression of associated proteins such as caveolin 3 (CAVIN3), caveolin 1 (CAV1), vimentin (VIM), cortactin (CTTN), cofilin 1 (CFL1), and Actinin Alpha 4 (ACTN4) (Table 1) (Sup. Fig. 2 B, Sup. Table 2). Moreover, these data suggest that DCLK1 might contribute to the enhanced motility of tumor cells.

**Figure 1.**
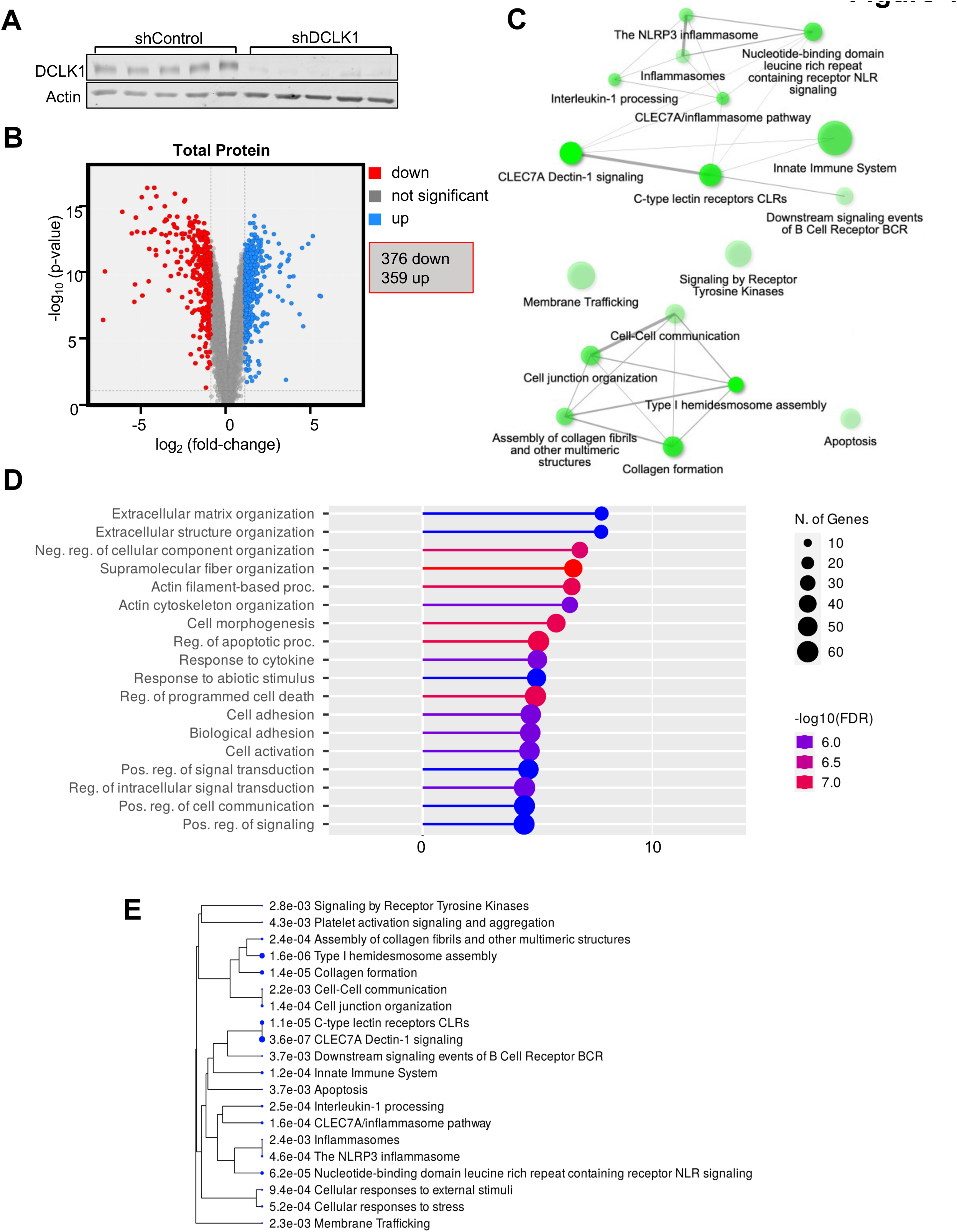
DCLK1 knockdown regulates expression of a number of genes. (A) Immunoblot analysis of DCLK1 knockdown in HNSCC FaDu cells is presented, with five independent experimental replicates confirming consistent reduction in DCLK1 levels compared to the control. Actin was used as a loading control to ensure equal protein loading across all lanes. (B) A volcano plot is provided, depicting proteins that are either downregulated (in red) or upregulated (in blue) in shDCLK1 cells relative to the control, with log2 fold change greater or less than 1.0. Functional annotation of DCLK1 is detailed, including gene ontology enrichment analysis of (C) pathway network enrichment, (D) biological processes, and (E) hierarchical dendrogram. Fisher’s exact test was employed for the functional annotation, with a significance threshold of P-value < 0.05 determined using the Benjamini-Hochberg procedure to identify statistically significant values. The figures were generated using ShinyGO.

To further understand the functional significance of DCLK1-mediated proteomic and phosphoproteomic changes, we conducted an integrative analysis to identify the regulatory kinases that may potentially be involved in mediating DCLK1’s cellular functions. We utilized curated libraries like BioGRID [19], MINT [20], STRING [21], and others, facilitated by the KEA3 web tool [22]. Notably, the analysis revealed reduction in the levels of several kinases in the DCLK1 knockdown cells, including Src kinase, epidermal growth factor receptor (EGFR), and P21 (RAC1)-activated kinase 1 (PAK1), aligning with our earlier gene ontology analysis emphasizing roles in locomotion and cytoskeletal rearrangement (Sup. Fig. 3A, Sup. Table 3).

The human kinome, composed of over 500 kinases grouped into distinct superfamilies, poses complexity for visualization. To address this challenge, we employed Coral, a quantitative kinase visualization tool [23]. Our analysis revealed significant effects of DCLK1 knockdown on various kinases, consistent with our previous findings. EGFR emerged as one of the prominently downregulated kinases in the DCLK1 knockdown cells. Additionally, the CAMK kinase family exhibited susceptibility to DCLK1 downmodulation, highlighting DCLK1’s significant role within this kinase family (Sup. Fig. 3 B).

Our phosphoproteomic analysis also revealed a notable distinction in phosphopeptides, with 824 unique peptides exhibiting higher levels in DCLK1 knockdown cells compared to control, while 1,249 unique peptides were significantly lower (Fig. 2A, Sup. Table 1). To further explore the functional implications, we examined kinase activity associated with DCLK1 using the RoKAI/RoKAI Explorer tool [24]. Additionally, we conducted a functional ontology analysis based on the comparison of total phosphopeptides to total peptides. This analysis highlighted top-level ontology terms linked to cytoskeletal rearrangement and movement, exemplified by actin filaments and microtubules. Simultaneously, it revealed a notable enrichment of terms associated with membrane interaction, encompassing plasma membrane localization, cell junctions, and focal adhesion, in the DCLK1-enriched sample. (Fig. 2B). These findings strongly implicate DCLK1 in cellular motility, invasion, and cytoskeleton interactions.

**Figure 2.**
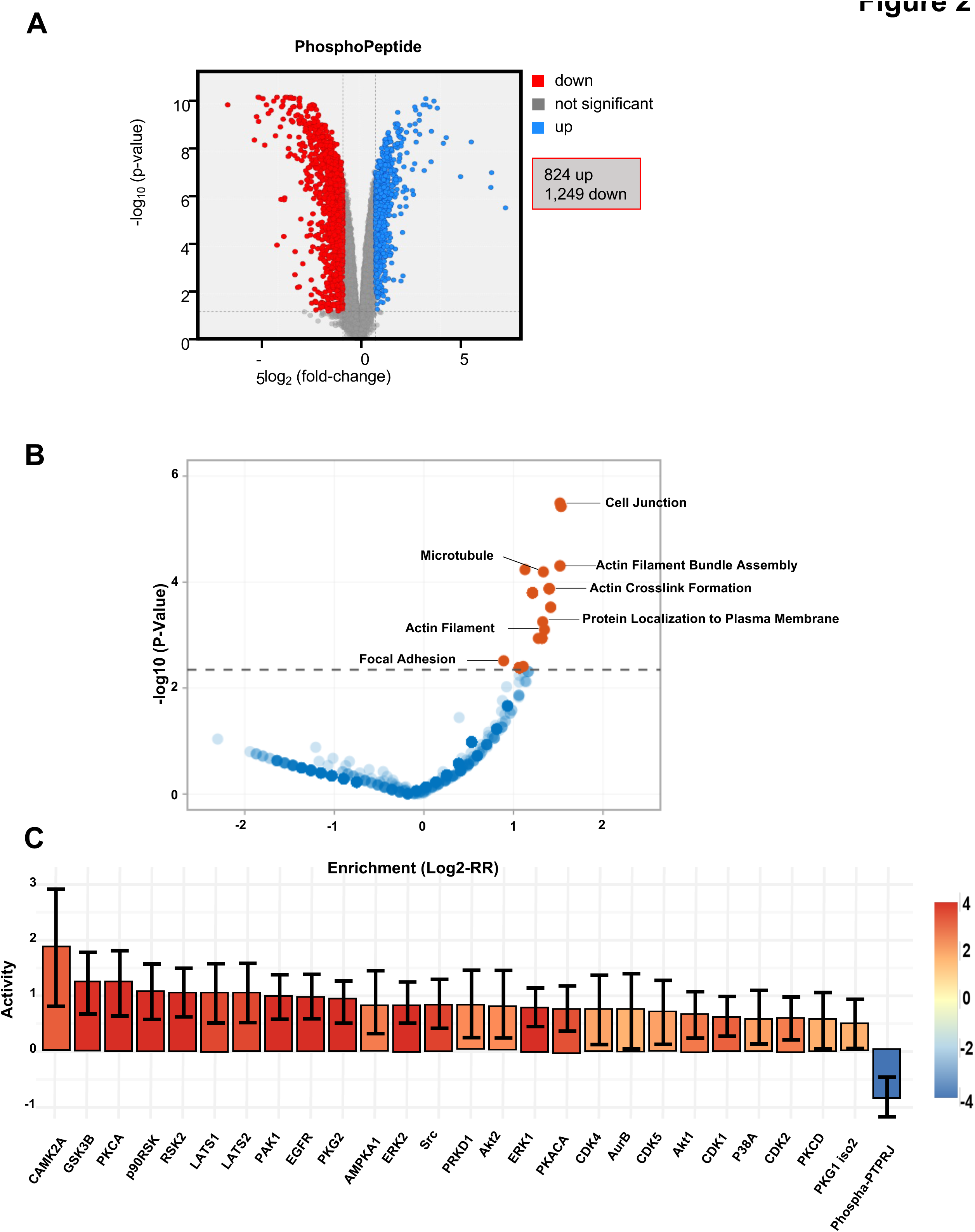
DCLK1 regulates the phosphoproteome. (A) A volcano plot is provided, depicting phosphopeptides that are either downregulated (in red) or upregulated (in blue) in shDCLK1 cells relative to the control, with log2 fold change greater or less than 1.0 FaDu cells. (B) The volcano plot showcases the outcomes of Gene Ontology analysis derived from inferred phospho-kinase analysis of phosphopeptides using the RoKAI Explorer web tool [24]. The dashed line indicates the significance threshold, with significant Gene Ontology terms highlighted in red and insignificant terms in blue. The maximum false discovery rate is set at 0.05, and the minimum absolute log2-fold change is set at 1.0. (C) The bar chart represents inferred phospho-kinase activity generated through the RoKAI web tool [24]. Each bar’s color corresponds to the z-score level of the kinase, with red indicating a higher score and blue indicating a lower score. Activity levels are determined based on the log2-fold change of phosphopeptides that are differentially expressed between shControl cells and shDCLK1 cells.

Through individual kinase activity assessment, our quantitative approach unveiled significant associations between DCLK1 and key kinases, including EGFR, ERK1/2, Src, and PAK1 (Fig. 2 C, Sup. Table 4). These findings provide additional insights into DCLK1’s impact on essential cellular functions, particularly its regulatory role in pathways associated with cell invasion.

### DCLK1 localizes to HNSCC invadopodia, driving cellular invasion

Unfavorable outcomes in HNSCC patients are often associated with locoregional invasion, a less explored aspect of the disease. Our previous work demonstrated that DCLK1 knockdown reduces cell invasion [15]. In invasive solid tumors, ECM degradation occurs through direct cell-to-matrix contact, facilitated by specialized subcellular invasive structures called invadopodia, with finger-like morphology [25]. In the mature invadopodia phase, microtubule extension is crucial for stabilization and cargo trafficking. The mechanism of MMP transport to the distal end of the invadopodia for ECM degradation has not been fully characterized. TCGA analysis further underscores DCLK1’s role by revealing its association with advanced HNSCC lymph node staging and higher tumor histologic grade (Fig. 3 A and B, respectively). Moreover, by analyzing immunohistochemical (IHC) specimens obtained from HNSCC patients (n = 17), we observed an elevated staining pattern concentrated specifically at the invasive boundary, where the tumor interacts with the surrounding stroma (Fig. 3 C). Given DCLK1’s function in trafficking of proteins in neuronal cells, we hypothesized that DCLK1 could facilitate HNSCC invasion by transporting MMP cargo to the distal end of lengthening invadopodia.

**Figure 3.**
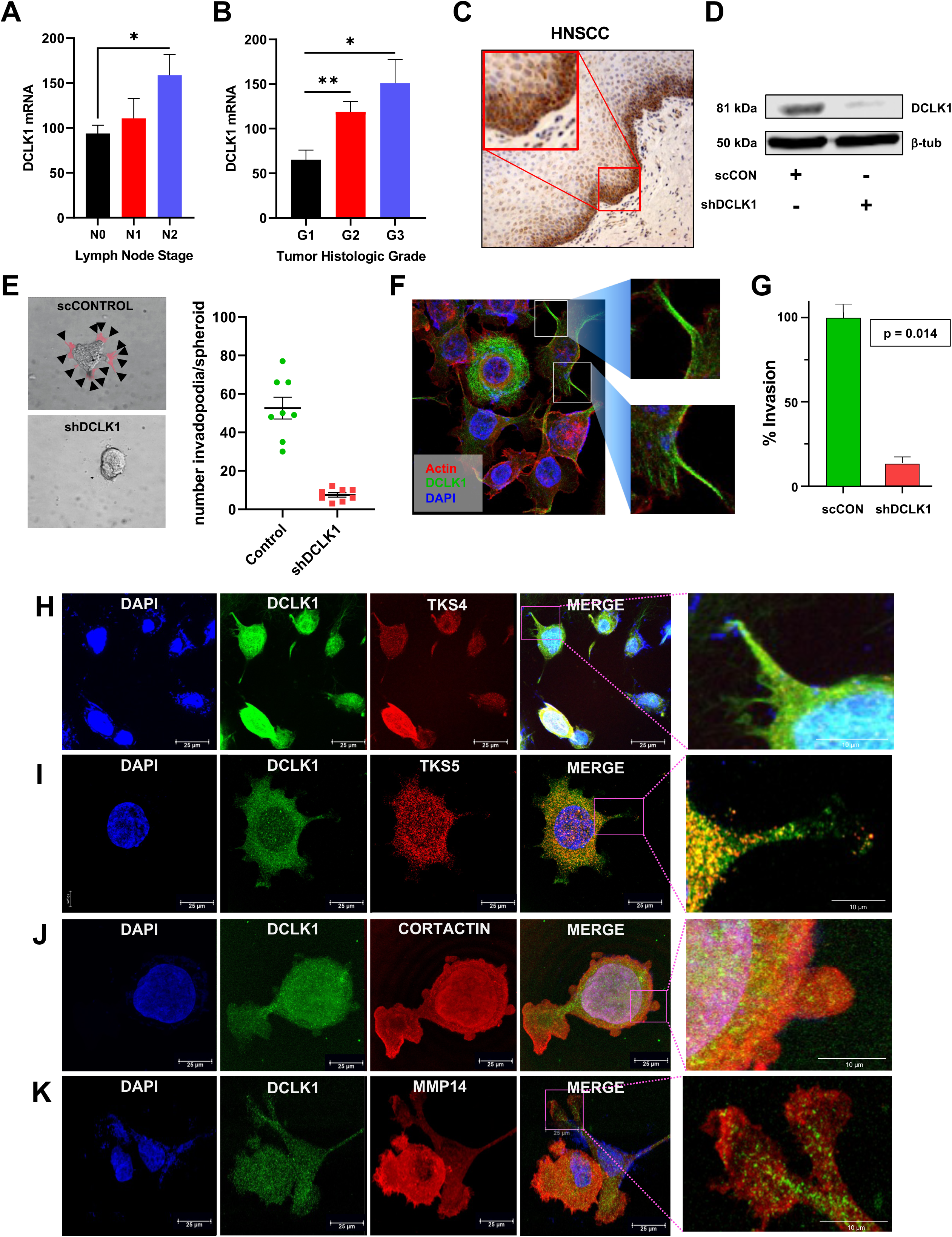
DCLK1 promotes HNSCC cellular invasion. Clinical data from TCGA reveal heightened DCLK1 expression in (A) progressed lymph node stage and (B) escalating histologic grade of head and neck squamous cell carcinoma (HNSCC). (C) Representative immunohistochemistry (IHC) staining of a HNSCC patient sample depicts elevated DCLK1 staining at the tumor-stroma interface (40x magnification, n = 27). (D) Successful knockdown of the 82 kDa DCLK1 protein in FaDu cells demonstrated by Western blot. (E) Abrogation of HNSCC protrusions (red colorization, black arrows) upon DCLK1 knockdown. Box and whisker plot quantifies invadopodia in 5 x 10^3^ number of cells across three replicate experiments using ImageJ. (F) Confocal images illustrate DCLK1 (green) localization in invadopodia of FaDu HNSCC cells, showcasing colocalization with actin (red) within invadopodia rather than the cytosol. Images captured using a 40X oil immersion lens objective. (G) Significant reduction in HNSCC cell invasion across the Boyden chamber, normalized to cell viabilities. Error bars indicate ± SEM. Confocal imaging of DCLK1 (green) with invadopodia markers (red) and DAPI reveals colocalization of DCLK1 with (H) TKS4, (I) TKS5, (J) cortactin, and (K) MMP14. Oil immersion at 100x objective.

Invadopodia play a pivotal functional role in initiating the degradation of the surrounding ECM barrier, and subsequently penetrating into the subsequent space created. We established an invadopodia assay wherein HNSCC cells were seeded onto a thin layer of VitroGel product. Following 30 minutes to allow for cell adhesion, additional matrix material was layered on top of the cells, creating a sandwich-like structure with the cells enclosed between layers of extracellular matrix (ECM). Previous reports have indicated varying maturation times for invadopodia, ranging from 3 minutes to 90 minutes [26–28].

Since targeting DCLK1 affects cell proliferation, we evaluated the effect of DCLK1 modulation on invadopodia prior to cell doubling. Since FaDu has a doubling time of 48 hours, cells were fixed and assessed using immunofluorescence 12 h post seeding. We observed a significant reduction in the number of cellular projections in DCLK1 knockdown cells compared to the wild-type control cells (Fig. 3 E).

To confirm the role of DCLK1 in invadopodia formation, we evaluated its localization within the cell using super-resolution microscopy. The data demonstrate that DCLK1 (green) localizes to HNSCC invadopodia (Fig. 3 F). Additionally, we observed actin (red) localization, consistent with established knowledge of the actin-rich nature of invadopodia [30]. Importantly, the localization patterns of DCLK1 and actin within invadopodia were not extensively overlapping (Fig. 3 F insets). The observed elongated, linear patterns of DCLK1, protruding parallel to cellular extensions, indicate that the localization of DCLK1 to invadopodia is not contingent upon actin-DCLK1 co-localization.

Invadopodia are the functional cellular units of invasive tumor cells; so, we conducted a Boyden chamber experiment to assess the impact of DCLK1 on the invasive potential of HNSCC cells. In this experiment, HNSCC cells were seeded onto a VitroGel substrate in the upper chamber and allowed to invade through the substrate towards the lower chamber. Our findings demonstrated that DCLK1 knockdown resulted in a significant reduction in the number of cells capable of effectively penetrating the gel and traversing to the lower chamber (p = 0.014). (Fig. 3 G).

To establish DCLK1 as a bonafide component of invadopodia, we conducted co-immunofluorescence studies with DCLK1 and several known markers that play distinct roles in invadopodia formation. The analysis revealed significant colocalization between DCLK1 and TKS4, an adapter protein predominantly present in the mature invadopodia (Fig. 3 H, Sup. Fig 4). Upon DCLK1 knockdown, we observed reduced fluorescence in knockout cells, coinciding with a decrease in TKS4 (Sup. Fig. 4 A). Proximity ligation assay (PLA) confirmed the close spatial interaction between DCLK1 and TKS4 (Sup. Fig. 4 B). Additionally, DCLK1 exhibited colocalization with TKS5, an adapter protein involved MMP transport (Fig. 3 I, Sup. Fig. 5 B). Furthermore, we observed colocalization between DCLK1 and cortactin, a key cytoskeleton organizer, indicating its involvement in cytoskeletal rearrangement during invadopodia formation (Fig. 3 J, Sup. Fig. 5 C). Lastly, DCLK1 and MMP14, a membrane-bound metalloprotease responsible for ECM degradation, were both seen in invadopodia (Fig. 3 K, Sup. Fig. 5 D).

The dynamics of the cytoskeleton play a pivotal role in governing both invadopodia dynamics and the adhesive capacity of cancer cells across diverse extracellular substrates [32]. Accordingly, we performed cell adhesion assays using collagen, fibronectin, vitronectin, and laminin substrates to evaluate the role of DCLK1 in cell adhesion. Our findings revealed that DCLK1 knockdown resulted in reduced cell adhesion to all substrates except fibronectin, indicating the involvement of DCLK1 in mediating cell-substrate interactions (Sup. Fig. 6 A and B).

### DCLK1 regulates the secretion of matrix metalloproteases

The presence of DCLK1 in invadopodia, particularly during the mature phase, is strongly supported by its co-localization with markers like TKS4 and cortactin. Functional studies involving DCLK1 knockdown have shown a reduced invasive capacity of HNSCC cells. Additionally, considering DCLK1’s association with cargo trafficking in neurons, further investigation into its potential influence on the secretion of degradative MMPs is warranted, shedding light on its role in ECM degradation and invasive behavior in HNSCC.

MMP2 and MMP9, known as gelatinases, possess a unique ability to break down gelatin, a protein derived from collagen, which is crucial for degrading ECM substrates. To detect MMP activity at the leading edge of the cell, we use dye-quenched (DQ)-gelatin, composed of quenched FITC-labeled gelatin. Upon gelatinolytic activity, the DQ substrate transforms into brightly fluorescent peptides, demonstrating gelatinase activity. When embedded in DQ-saturated VitroGel, our observations indicated a significant reduction in gelatin degradation after DCLK1 knockdown, strongly implying that DCLK1 actively promotes the secretion of gelatinases (Fig. 4 A). We further utilized the DQ assay to evaluate areas of degradation that overlap with HNSCC cells seeded atop the gel. We found that the localization of DCLK1 on the ventral surface of the cell concomitated with focal degradation of gelatin (Sup. Fig. 7).

**Figure 4.**
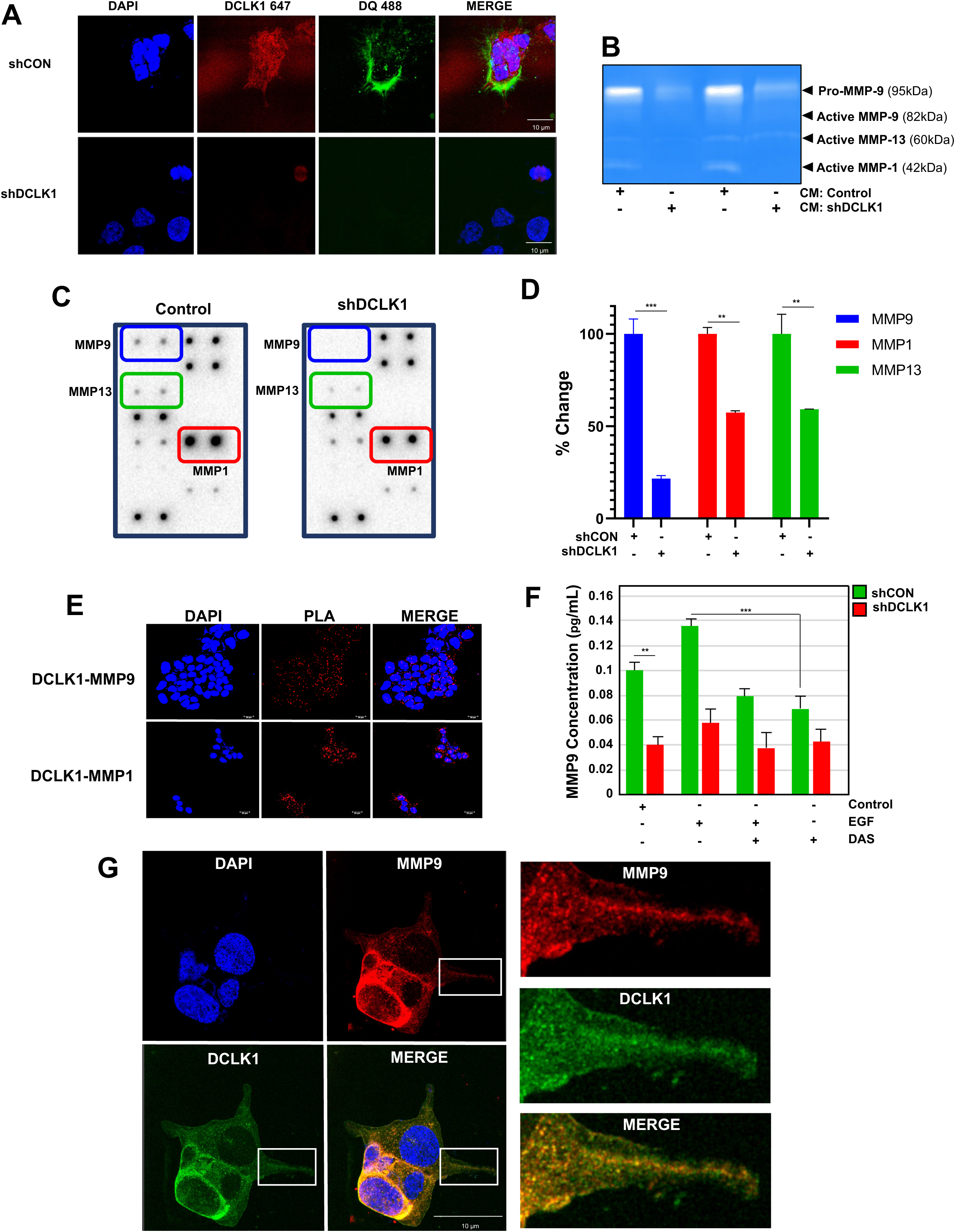
HNSCC invasion and cellular secretion of MMPs are impaired by DCLK1 knockdown. (A) Representative confocal imaging depicts DCLK1 (red) along with DQ degradation (green) in FaDu cells. Notably, DCLK1 knockdown leads to a marked reduction in DQ gelatin degradation. (B) Gelatin zymography reveals concentrated supernatants from shControl or shDCLK1 cells. White bands against a stained blue background indicate degradation areas, with specific MMPs identified based on their known molecular weights. (C, D) Conditioned media supernatants from shControl or shDCLK1 cells were analyzed using an MMP array. Quantification highlights significant differences in MMP levels between the two conditioned media groups. Error bars represent ± SEM. (E) Confocal imaging of PLA between DCLK1-MMP9 or DCLK1-MMP1 in FaDu HNSCC cells reveals close proximity, indicating a tight association. Images obtained at 40x oil immersion objective (representative image of 3 biological replicates from FaDu cells. (F) MMP9 ELISA bar graph compares conditioned media from shControl (green) and shDCLK1 (red) under different treatment conditions. Error bars represent ± SEM. ** P<0.001, *** P<0.0001 (G) Super resolution confocal microscopy demonstrate DCLK1 (green) and MMP9 (red) colocalize (yellow) specifically at the invadopodia of FaDu cells. Images on the right are digital magnifications of the inset white box in the 100X oil immersion objective image in FaDu cells.

To examine the impact of DCLK1 knockdown on MMP secretion we interrogated conditioned media (CM) from shControl and shDCLK1 HNSCC cells. Initially, we conducted zymography using SDS page gels embedded with gelatin to visualize and quantify enzyme activity in the CM samples. Intriguingly, after DCLK1 knockdown, we observed reduced secretion of both pro and active forms of MMP9, active MMP13, and MMP1. Notably, MMP2 secretion was not significantly detected in either the knockdown or control CM samples (Fig. 4 B). To further validate and quantify MMP secretion levels in CM, we employed an MMP array. Our results confirmed reduce levels of MMP13 and MMP1 in CM from DCLK1 knockdown HNSCC. Additionally, DCLK1 knockdown led to a significant decrease in MMP9 secretion from HNSCC (p < 0.001) (Fig. 4 C, D). Employing PLA, we demonstrate that DCLK1 showed proximity with both MMP9 and MMP1, with a higher frequency of interactions observed between DCLK1 and MMP9 (Fig. 4 E). This further demonstrates that DCLK1 associates with MMPs, particularly with MMP9.

EGFR signaling leads to c-Src activation, subsequently promoting actin assembly, polymerization, and cortactin activation, which results in invadopodia formation [33]. Dasatinib, a potent Src inhibitor, effectively inhibits Src activity [34]. To investigate the impact of DCLK1-mediated MMP9 secretion, we conducted an enzyme-linked immunosorbent assay (ELISA) using CM from HNSCC cells. Initially, when HNSCC cells were exposed to EGF, we observed an increase in MMP9 secretion, indicating that EGF promotes MMP9 secretion (Fig. 4 F). Conversely, treatment with Src inhibitor, dasatinib resulted in a noticeable reduction in EGF induced MMP9 secretion. MMP9 levels were also decreased in the shDCLK1 group compared to the control. Treatment with dasatinib did not result in a further reduction in MMP9 levels, underscoring the significance of DCLK1 in mediating MMP9 secretion. Likewise, gel zymography validated an overall dampening of shDCLK1 CM to degrade gelatin, despite EGF stimulation (Sup. Fig. 8 A)

We sought to determine if DCLK1 actively participated in the intracellular trafficking of MMPs, to the distal end of invadopodia using confocal microscopy. Our findings revealed a clear colocalization between DCLK1 and MMP9, within invadopodia. Notably, DCLK1 displayed well-organized, centrally located extensions that reached towards the distal ends of invadopodia (Fig. 4 G). This spatial arrangement strongly suggested that DCLK1 actively regulated MMP secretion within the context of invadopodia, contributing to their role in promoting invasive behavior.

In all our experimental analyses, including zymography and MMP array assays, we consistently observed that DCLK1-mediated MMP secretion did not involve MMP2. To confirm this, we conducted confocal imaging and conducted co-staining for DCLK1 and MMP2. These results further supported the findings from the MMP array, as we observed minimal colocalization between DCLK1 and MMP2 (Sup. Fig. 8 B). These results indicate DCLK1 most closely interacts with MMP 1 and 9 and not MMP2.

### DCLK1 regulates the KIF16B motor protein

MMP secretion is a critical function of mature and activated invadopodia [35]. Given that MMPs are transported by motor proteins, our goal was to identify the specific motor proteins interacting with DCLK1. To accomplish this, we conducted an extensive screening of pan-kinesins utilizing TCGA. We observed a strong correlation between DCLK1 and the Kinesin 3 sub-family across all stages of HNSCC (Fig. 5 A). Within this group, KIF1B exhibited high levels of correlation with DCLK1, particularly as the tumor stage increased. Additionally, ChipSeq experiments assessed using TNMplot [36] demonstrate correlation of DCLK1 to KIF1A, KIF1B, KIF1C, and KIF16B compared to normal head and neck tissue (Fig. 5 B). Likewise, gene expression of the Kinesin-3 family is starkly different from normal, demonstrating a dramatic increase in KIF1B, KIF1A, KIF14, and especially KIF16B (Fig. 5 C).

**Figure 5.**
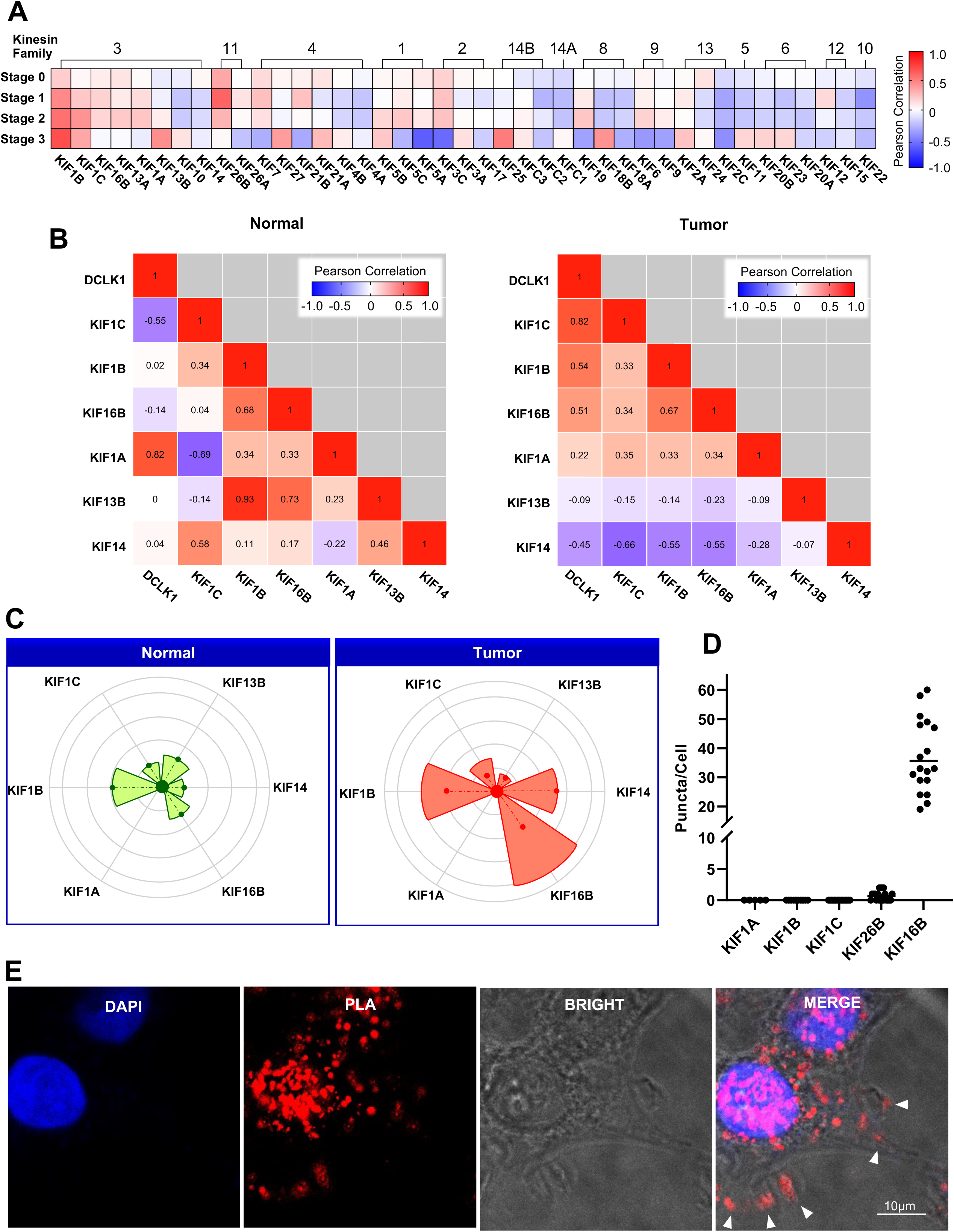
DCLK1 is associated with the kinesin 3 motor protein KIF16B. (A) Heatmap displaying the correlation between DCLK1 and members of the kinesin superfamily in various stages (0-3) of HNSCC. Notably, members of the kinesin 3 family exhibit the most substantial positive correlation with DCLK1. (B) Correlation matrices from ChIP-seq data illustrate the diverse patterns of kinesin 3 family members in normal and HNSCC tumors, influenced by DCLK1 expression. (C) Protein expression analysis reveals distinct patterns of kinesin 3 family members in tumors compared to normal tissue, with the pie size indicating fold change and concentric circles representing varying p-value thresholds for statistical significance. (D) Bar graph depicting the results of PLA screen, indicating the proximity of DCLK1 to different kinesin 3 family members. Error bars represent ± SEM. (E) Confocal imaging of PLA between DCLK1 and KIF16B in FaDu demonstrates a substantial interaction, evident from numerous PLA puncta. Images acquired using a 40x oil immersion objective.

Next, we utilized the PLA technique to identify the specific members of the Kinesin 3 family that are in proximity with DCLK1. Surprisingly, our results showed that none of the previously reported Kinesin 3 family members, known for their strong interactions, exhibited a discernible preferential association with DCLK1 in HNSCC cells (Fig. 5 D; Sup. Fig. 9). Although existing literature has hinted at DCLK1’s association with the Kinesin 3 subfamily, as indicated in the pan-kinesin TCGA screen (Fig. 5 A), most studies have suggested an interaction between DCLK1 and KIF1A in neuronal cells [17]. However, using PLA we demonstrate a proximity between DCLK1 and KIF16B, an anterograde kinesin (Fig. 5 E and Sup. Fig. 10, respectively). Our findings indicate a distinct preference of DCLK1 for binding with KIF16B. To validate the colocalization of DCLK1, TKS4, and KIF16B, we conducted immunofluorescence studies using HNSCC cell lines treated with EGF (5 min), dasatinib (2 hr), a combination of dasatinib and EGF, or a DMSO control. Our results demonstrated that EGF stimulation increased the colocalization of DCLK1, KIF16B, and TKS4 (Sup. Fig. 11). Conversely, the inhibition of Src using dasatinib disrupted the colocalization with DCLK1.

### DCLK1, KIF16B, RAB40B form a complex

Following the confirmation of the binding interaction between DCLK1 and KIF16B and their potential role in transporting MMP cargo, our next aim was to provide evidence of tight proximity with spatial resolution inside HNSCC. We performed PLA for DCLK1, KIF16B, and RAB40B and observed that even in controls, there were a few complexes between the three proteins within the cell, which was further enhanced following EGF treatment (Fig 6 A, Sup. Fig 12). However, this interaction was suppressed when cells were also treated with dasatinib (Fig 6 A and B). Furthermore, *in-silico* modeling analysis confirmed the binding interaction between the DCLK1 kinase domain and the motor head domain of KIF16B. Specifically, the binding occurs at arginine 442 in DCLK1 and aspartic acid 1199 in KIF16B, with a distance of 2.6 angstroms (Sup. Fig 13).

**Figure 6.**
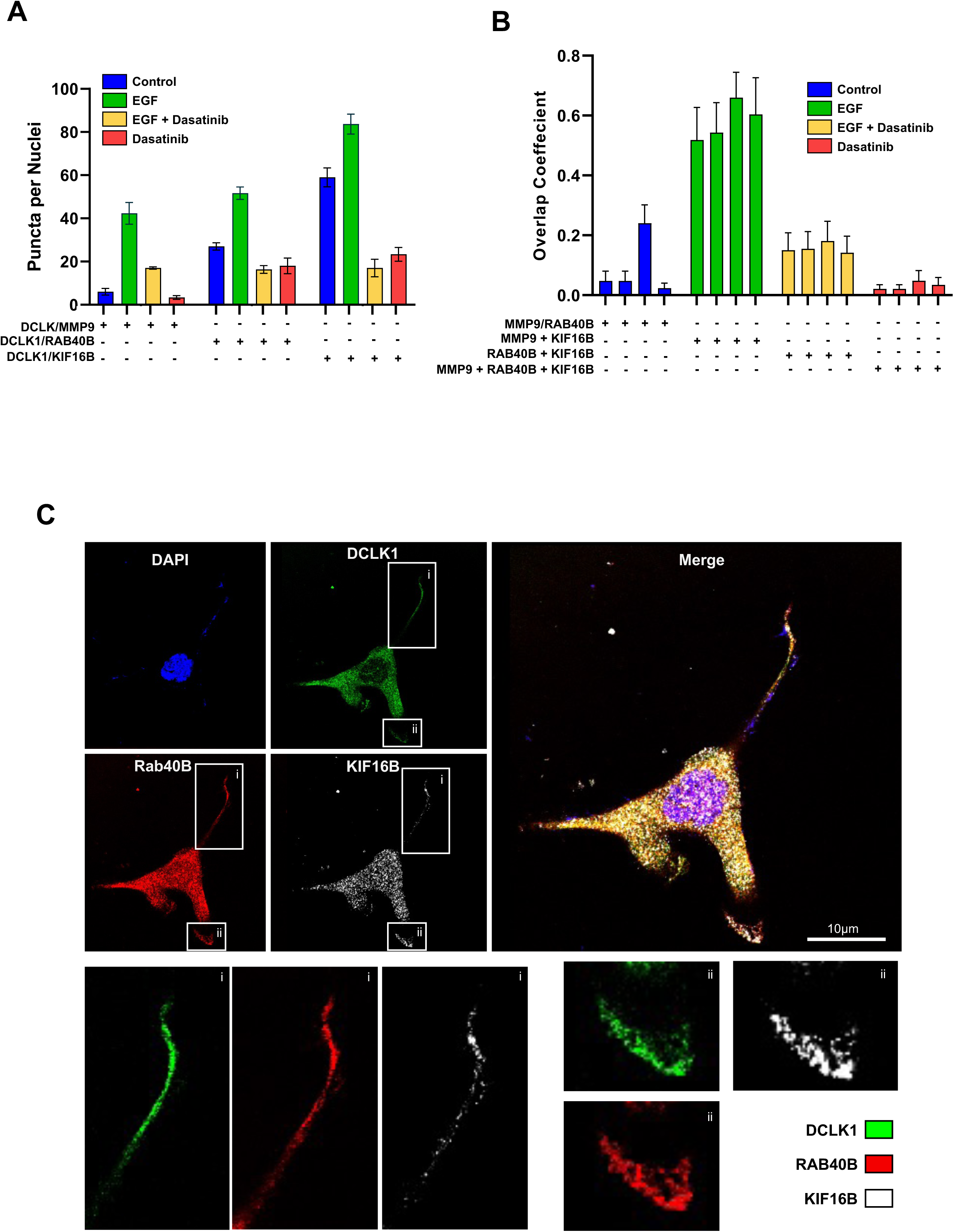
DCLK1, KIF16B, and RAB40B form a complex in HNSCC. (A) Quantification of proximity ligation assay (PLA) results depicting the association of DCLK1 to MMP9, RAB40B, and KIF16B in UMSCC1 HNSCC cells under different conditions: control (blue), EGF (10 ng/ml) stimulation (green), EGF and dasatinib (100nM) treatment (yellow), or dasatinib alone (red). The bar graph illustrates the changes in PLA signals. Error bars indicate ± SEM. The graph is a representative image from three biological replicates conducted in FaDu, UMSCC1, and CAL33 cells. (B) Quantification of three-color PLA demonstrating overlap coefficient analysis showing association between DCLK1 and each of MMP9, RAB40B, or KIF16B under the experimental conditions mentioned in 6A. Error bars represent ± SEM. (C) Confocal microscopy images capturing the localization of DCLK1 (green), Rab40B (red), and KIF16B (white) in invadopodia of FaDu cells. The merged image highlights significant colocalization of these proteins within invadopodia. Each inset (i, ii) showcases individual cellular projections. Images were obtained using oil immersion at a 100x objective.

Additionally, KIF16B, RAB40B, and DCLK1 colocalize within invadopodia (Fig. 6 C). Using confocal imaging, we observed an overlap among DCLK1, KIF16B, and RAB40B, with all three proteins localizing to invadopodia. This observation solidifies the presence of a complex within invadopodia, composed of DCLK1, KIF16B, and RAB40B, collectively contributing to the transport of degradative cargo in these specialized structures.

## Discussion

Invadopodia-mediated degradation plays a critical role in promoting locoregional invasion in HNSCC, significantly impacting overall mortality [37]. While extensive research has focused on understanding the activation and initiation of invadopodia, particularly their extracellular to intracellular dynamics, our knowledge of the mature phase of invadopodia, involving microtubule extension and cargo transportation in HNSCC, remains limited [38]. Although it’s clear that vesicle transfer occurs along invadopodia in a plus-end direction facilitated by microtubules, investigations into the specific regulatory factors in this process are lacking [35]. In this context, DCLK1 stands out as a compelling candidate due to its potential role as a microtubule binding protein known for participating in cargo transport and driving cancer progression [17].

Previously, DCLK1’s role in promoting cancer cell invasion has been predominantly attributed to its facilitation of EMT, with limited attention given to its direct association with the invasion machinery [39]. Earlier studies utilizing Boyden chamber assays focused solely on invasion function, yielding relatively binary results indicating the presence of cells that have invaded through a barrier without elucidating the mechanism of invasion. To address this knowledge gap, our research replicated trans-well experiments and specifically examined the presence of DCLK1 within invadopodia. Invadopodia are specialized structures primarily responsible for secreting degradative proteinases during the mature phase of invadopodia formation [40]. While previous studies on MMP secretion in HNSCC mainly focused on vesicular trafficking, particularly targeting to the plasma membrane, the specific kinesins responsible for MMP trafficking in HNSCC and the role of kinesin binding elements remained unknown [35, 40].

In our investigation, we explored the impact of DCLK1 knockout on MMP secretion in HNSCC CM, with a particular focus on gelatinase MMP9, given its relevance in ECM degradation and overexpression in HNSCC [41]. Intriguingly, we observed a significant reduction in MMP secretion, especially MMP9, in CM from DCLK1 knockout cells. This suggests for the first time, a critical role for DCLK1 in regulating MMP9 secretion within invadopodia. Importantly, we also observed gelatinase activity in DCLK1-mediated secretion, further supporting the connection between DCLK1 and MMP9 in ECM degradation during cellular invasion and local metastasis.

On the other hand, MMP2, with its broader functional role, is primarily associated with angiogenesis, a distinct process separate from the ECM degradation orchestrated by invadopodia [42]. In contrast, MMP9 displays exceptional degradative activity against the most prevalent ECM substrates found in HNSCC tumors, such as collagen III, IV, and V [43].

While RAB40B has been linked to the vesicular transport of matrix metalloproteases within invadopodia in breast cancer, the specific mechanism governing its locomotion remains to be elucidated [13, 35]. On the other hand, KIF16B features a highly processive motor domain that allows for rapid movements up to 10 µm [44]. This motor protein is instrumental in recycling MT1-MMP from the plasma membrane, thereby facilitating macrophage invasion [45].

Furthermore, the interplay between DCLK1 and the kinesin 3 superfamily, which encompasses KIF16B, is responsible for directing dense core vesicles towards the microtubule plus end [17]. Our research represents a groundbreaking contribution by being the first study to explore the interaction between the microtubule-binding protein DCLK1 and the functional role of KIF16B within the cancer milieu. Additionally, we test a hypothesis regarding the transport of degradative cargo by KIF16B in HNSCC. Finally, our work establishes a seminal connection, previously unidentified, among DCLK1, KIF16B, MMP9, and the RAB40B degradative complex in the context of cancer biology.

## Materials and Methods

### 3.1 Cells and Reagents

In this study, well-defined cell lines derived from HNSCC were utilized. The cell lines used were FaDu (ATCC), HN5 (obtained from Dr. Jeffrey Myers at The University of Texas MD Anderson Cancer Center), Cal33 (cat# ACC 447 DSMZ; Braunschweig, Germany), and UMSCC1 (obtained from Dr. Thomas E Carey at the University of Michigan). The authentication of established cell lines was performed periodically at Johns Hopkins utilizing the Promega GenePrint 10 kit, and the analysis was conducted utilizing GeneMapper v4.0. Each cell line was cultured in DMEM (Corning) supplemented with 10% heat-inactivated FBS (Sigma-Aldrich) not including antibiotics. Incubation of cells was carried out at 37°C with 5% CO2. Tissue Microarray (#HN483) was acquired from US Biomax, Inc. The following antibodies were used: DCLK1 (ab31704), TKS4 (ab122342), MMP9 (ab73734), MMP2 (ab92536), MMP1 (ab52631), MMP13 (ab219620) from abcam (Cambridge, UK); TKS5 (NBP1-90454), KIF16B (NBP1-14157), KIF1A (NBP1-80033), KIF1B (NB100-57493), KIF1C (NBP1-85978), KIF26B (NBP1-90443) from Novus Biologicals (Centennial, CO). β-tubulin from Sigma-Aldrich; pan-actin (H0000059-K) from Abnova (Taipei, Taiwan); anti-rabbit IgG Dylight 680 (#35568), anti-rabbit IgG Dylight 488 (#35553), and anti-mouse IgG Dylight 800 (#35521) from ThermoFisher (Waltham, MA). Texas Red-X Phalloidin (T7471) and DAPI 1306 were used for nuclear counterstaining (ThermoFisher). VitroGel Hydrogel Matrix-3D Cell Culture (VHMO1 and TWG001) from The Well Bioscience (North Brunswick, NJ) was employed. The N-Terminal pFLAG 3 vector (Addgene) was employed for cloning the DCLK1 DCX DNA by restriction digest, utilizing NotI and BamHI restriction enzymes. Following transfection, FaDu HNSCC cell line was isolated using G418 sulfate salt at a concentration of 400 µg/ml. In relation to the DCX Domain, the primers utilized for DCLK1 NotI were as such: Forward Primer (FP): 5′-GATCGCGGCCGCGATGTCCTTCGGCAGA-3′, and Reverse Primer (RP): 5′-GATCGGATCCCTAGCCATCGTTCTCATC-3′. For tandem mass tagged isobaric labeling, the TMT10plex (90110) kit from ThermoFisher was employed.

### 3.2 Immunoblot

Cellular proteins were obtained by extracting whole-cell lysates using RIPA lysis buffer enhanced by addition of phosphatase and protease inhibitors (Roche 11836153001). To remove debris, the cells were sonicated while on cold blocks and centrifuged, and the resulting supernatants were kept at -80°C. Subsequently, the protein from cells were parted by electrophoresis on 10% SDS-polyacrylamide gels and moved onto PVDF membranes. To prevent nonspecific binding, the membranes were blocked using a mixture of TBS Intercept blocking buffer (Licor 927-60050) and TBS in a 1:1 ratio. Primary antibodies, diluted in a 1:1 blocking buffer to TBST, were then incubated overnight with the membranes. Banding from protein was visualized using the LI-COR Odyssey DLx method and quantified utilizing ImageJ.

### 3.3 Mass Spectrometry

Total protein from each cell pellet was reduced, alkylated, and purified by chloroform/methanol extraction prior to digestion with sequencing grade modified porcine trypsin (Promega). Tryptic peptides were labeled using a tandem mass tag 10-plex isobaric label reagent set (Thermo) and enriched using High-Select TiO2 and Fe-NTA phosphopeptide enrichment kits in succession (Thermo) following the manufacturer’s instructions. Both enriched and un-enriched labeled peptides were separated into 46 fractions on a 100 x 1.0 mm Acquity BEH C18 column (Waters) using an UltiMate 3000 UHPLC system (Thermo) with a 40 min gradient from 99:1 to 60:40 buffer A:B ratio under basic pH conditions, and then consolidated into 18 super-fractions. Each super-fraction was then further separated by reverse phase XSelect CSH C18 2.5 um resin (Waters) on an in-line 150 x 0.075 mm column using an UltiMate 3000 RSLCnano system (Thermo). Peptides were eluted using a 60 min gradient from 98:2 to 60:40 buffer A:B ratio.Eluted peptides were ionized by electrospray (2.4 kV) followed by mass spectrometric analysis on an Orbitrap Eclipse Tribrid mass spectrometer (Thermo) using multi-notch MS3 parameters. MS data were acquired using the FTMS analyzer in top-speed profile mode at a resolution of 120,000 over a range of 375 to 1500 m/z. Following CID activation with normalized collision energy of 31.0, MS/MS data were acquired using the ion trap analyzer in centroid mode and normal mass range. Using synchronous precursor selection, up to 10 MS/MS precursors were selected for HCD activation with normalized collision energy of 55.0, followed by acquisition of MS3 reporter ion data using the FTMS analyzer in profile mode at a resolution of 50,000 over a range of 100-500 m/z.Buffer A = 0.1% formic acid, 0.5% acetonitrile;Buffer B = 0.1% formic acid, 99.9% acetonitrile.Both buffers adjusted to pH 10 with ammonium hydroxide for offline separation

### 3.4 Data Analysis – ProteoViz (phosphoTMT)

Proteins were identified and reporter ions quantified by searching the UniprotKB database restricted to Homo sapiens (January 2022) using MaxQuant (Max Planck Institute, version 2.0.3.0) [46]with a parent ion tolerance of 3 ppm, a fragment ion tolerance of 0.5 Da, a reporter ion tolerance of 0.001 Da, trypsin/P enzyme with 2 missed cleavages, variable modifications including oxidation on M, Acetyl on Protein N-term, and phosphorylation on STY, and fixed modification of Carbamidomethyl on C. Protein identifications were accepted if they could be established with less than 1.0% false discovery. Proteins identified only by modified peptides were removed. Protein probabilities were assigned by the Protein Prophet algorithm [47]. TMT MS3 reporter ion intensity values are analyzed for changes in total protein using the unenriched lysate sample. Phospho (STY) modifications were identified using the samples enriched for phosphorylated peptides. The enriched and un-enriched samples are multiplexed using two TMT10-plex batches, one for the enriched and one for the un-enriched samples.

Following data acquisition and database search, the MS3 reporter ion intensities were normalized using ProteiNorm [48]. The data was normalized using VSN [49] and analyzed using proteoDA to perform statistical analysis using Linear Models for Microarray Data (limma) with empirical Bayes (eBayes) smoothing to the standard errors [50]. A similar approach is used for differential analysis of the phosphopeptides, with the addition of a few steps. The phospho sites were filtered to retain only peptides with a localization probability > 75%, filter peptides with zero values, and log2 transformed. Limma was also used for differential analysis. Proteins and phosphopeptides with an FDR-adjusted p-value < 0.05 and an absolute fold change > 2 were considered significant.

### 3.5 Proteomic and Phospho-proteomic Analysis

An established cutoff of log2-fold change greater than 1.0 with a p-value <0.05 was used. To evaluate biological processes, gene ontology, and pathway networks associated with total protein data, the ShinyGO tool was employed.[51]. Top-level enriched gene ontology terms and biological pathways were determined by inputting differential proteins into Metascape [18]. Kinase enrichment analysis was performed using KEA3 [22]. The CORAL tool was used to establish the kinome tree and quantitatively evaluate kinase interactions [23]. For determining kinase activity through functional networks, the RoKAI app was utilized, and the beta version of RoKAI Explorer was employed to determine the functional ontology related to phosphor-kinases [24].

### 3.6 Invadopodium Assay

An angiogenesis chamber slide (Ibidi 81506) was prepared by allowing 10 µL of VitroGel (Company) to polymerize in the innermost well. Cells were subsequently seeded at 50% confluence and given less than 30 minutes for seeding. To create a hydrogel sandwich, 30 µL of VitroGel was added to the uppermost part of the well. The cell/VitroGel sandwich was then fixed with 4% PFA for staining, enabling confocal imaging, or viewed using bright field microscopy under live cell conditions.

### 3.7 Boyden Chamber Invasion Assay

Cell invasion was evaluated using the Transwell Boyden chamber system, following the protocol described by New et al. in 2017. In brief, 8-μm pore inserts were utilized for migration assessment. A thin layer of VitroGel was applied to the inserts. HNSCC cells (3 x 10^4^ cells/well) within serum-free culture media were plated on the VitroGel-coated inserts. These inserts were then set in triplicate containment wells containing media containing serum up to 24 hours. Simultaneously, cells (2 x 10^3^ cells/well in a 96-well plate) were plated under identical conditions to assess viability using the CyQUANT assay. Following fixative treatment then staining with the Hema 3 kit (Thermo Fisher Scientific), quantity of cells that invaded through the VitroGel and across the Boyden chamber membrane was quantified. The number of invading cells was normalized based on cell viability. To measure viability, HNSCC cells were seeded in quadruplicate (2 x 10^3^ cells/well in a 96-well plate) at initial day 0. Experimental conditions were applied on day 1 for durations of 24 to 72 hours, and cellular viability was quantified using a CyQUANT proliferation kit (Invitrogen C7026) pursuant to the manufacturer’s guidelines.

### 3.8 Cell Adhesion Assay

A total of 5 x 10^4^ cells per dish were seeded onto 60 mm dishes coated with fibronectin, type I collagen, laminin, or vitronectin. The cells were allowed to adhere for specific predetermined time points, followed by gentle washing with flowing PBS at 37°C for 20 seconds. Subsequently, the plates were fixed using 10% formalin (Fisher Scientific SF98-4) and stained with crystal violet. Images of three random fields were captured. The number of cell adhesion events and projections per cell were determined and quantified using ImageJ software (v1.53t).

### 3.9 Colony Formation Assay

HNSCC cells (3 x 10^3^ cells/well) were seeded in 6-well plates and permitted to grow until distinct, separate colonies became visible (21 days). The culture media was removed, and the cells were affixed using 10% formalin and stained via crystal violet. Scanning of the entire plates was performed, and the quantification of colony-forming units (CFUs) was conducted using ImageJ software (version 1.54).

### 3.10 Conditioned Media

HNSCC cells (5 x 10^7^ cells/well in a 145-mm dish), with or without DCLK1 knock out, were seeded in 10% FBS DMEM and cultured for 48 hours. Subsequently, the media was exchanged to non-serum containing DMEM for an additional 48 hours. During this period, 30 mL of fresh media was added every 24 hours. The media was then collected and concentrated using the Thermo Scientific Pierce 88527 kit, and the protein concentration was determined using the Bradford colorimetric assay.

### 3.11 Gel Zymography

Gel zymography analysis was conducted using 10% Zymogram gels (Novex ZY00100BOX) following the manufacturer’s instructions. In summary, 50 µg of CM from shDCLK FaDu or shControl FaDu cells was subjected to electrophoresis at a constant voltage of 125 V for 90 minutes. Following gel renaturation, the gel was incubated overnight and subsequently stained using SimplyBlue Safestain (Thermo LC6060). Clear bands on the gel indicate areas of protease activity, while the background remains blue.

### 3.12 Dye Quenched Assay

A volume of 10 µL of VitroGel, saturated with 50 μg/ml of DQ gelatin from pigskin (Thermo D12054), was polymerized in the central well of an angiogenesis chamber slide (Ibidi 81506). Subsequently, gel coverslips were prepared by seeding 2 × 10^5^ cells and incubating them at 37°C to allow degradation. Following incubation, the cells were fixed using 4% paraformaldehyde in PBS. Quantification of matrix degradation was performed by assessing the number of degraded spaces in 15 cellular fields (300-500 cells) for every experimental condition.

### 3.13 Proximity Ligation Assay

Cells were seeded in VitroGel at 50% confluence on chamber slides for PLA (Proximity Ligation Assay) using the Duolink *in situ* Detection Reagents Red kit (DUO92008, Sigma-Aldrich), following the given protocol. Various groupings of primary antibodies targeting DCLK1 and KIF16B, KIF1A, KIF1B, KIF1C, KIF26B, MMP1, MMP9, or RAB40B were employed. Subsequently, PLA puncta were imaged utilizing a TCS SP8 confocal laser scanning microscope (Leica Microsystems) equipped with a 40×1.35 NA or 100×1.35 NA oil immersion objective. High-resolution z-stack photograms were captured with a z-interval of 0.5 μm. Two-dimensional projections were generated using the LAX software platform (Leica Microsystems) by combining the maximum intensity from each z-series. The density of PLA puncta was determined by analyzing high-resolution images (captured with a 40×1.35 NA objective) using ImageJ (NIH).

### 3.14 Human MMP Antibody Array

The evaluation of human MMPs (Matrix Metalloproteinases) and TIMPs (Tissue Inhibitors of Metalloproteinases) was conducted following the instructions provided by the manufacturer (abcam 134004). In summary, 200 µg of total protein derived from CM of shDCLK1 or shControl FaDu cells was applied to separate blot arrays and incubated overnight. The arrays were subsequently subjected to chemiluminescence detection, and the resulting signals were captured at different exposure times.

### 3.15 The Cancer Genome Atlas Data Analysis

The gene expression RNAseq data of the TCGA head and neck cancer (HNSC) Firehose Legacy cohort was obtained using cBioPortal [52, 53]. The expression levels of DCLK1 were categorized as elevated to suppressed based on their relationship to the median quantification of gene-level transcripts estimated, which were converted using log2(x+1) and normalized using RSEM counts. These expression levels were then matched with clinical lymph node staging data obtained from TCGA.

### 3.16 Statistical analysis

The data are presented as the mean ± standard error of the mean (SEM). Significance in all experiments was assessed using non-parametric two-tailed Mann-Whitney U tests, and for comparison of multiple groups, the Kruskal-Wallis test was used. All statistical calculations were conducted via GraphPad Prism Software (version 9.1.0), and significance was decided by a p-value less than 0.05.

## Acknowledgement

This study was supported in part by the University of Kansas Cancer Center under CCSG P30CA168524; IDeA National Resource for Quantitative Proteomics and NIH/NIGMS grant R24GM137786.

The Leica STED microscope is supported by NIH S10OD023625 at the University of Kansas Medical Center, Kansas City, KS 66160.

Biomedical Research Training Program at the University of Kansas Medical Center supported graduate researcher stipend expenses during the formation of this work.

---

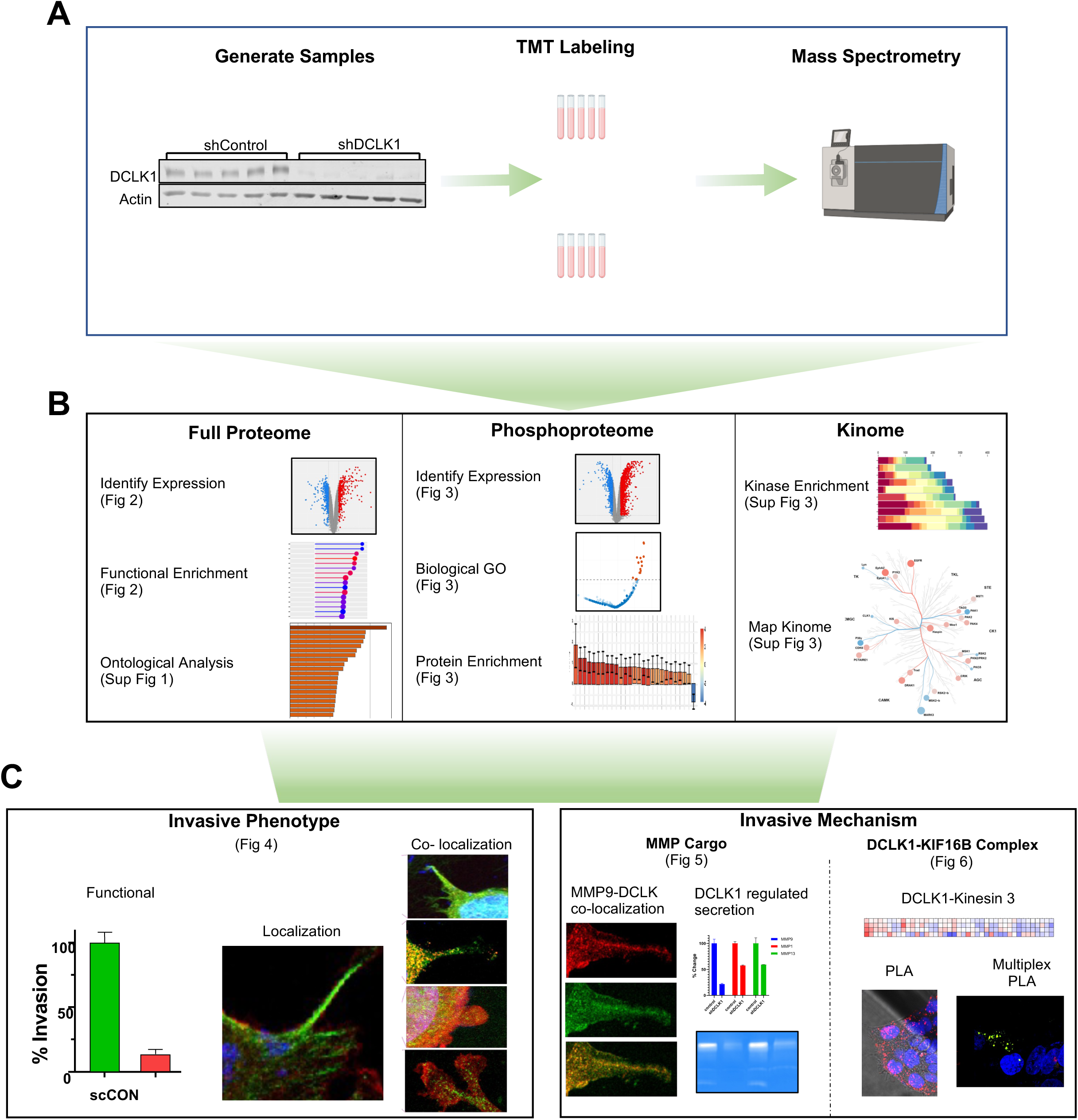
Graphical Abstract

**Supplemental Figure 1.**
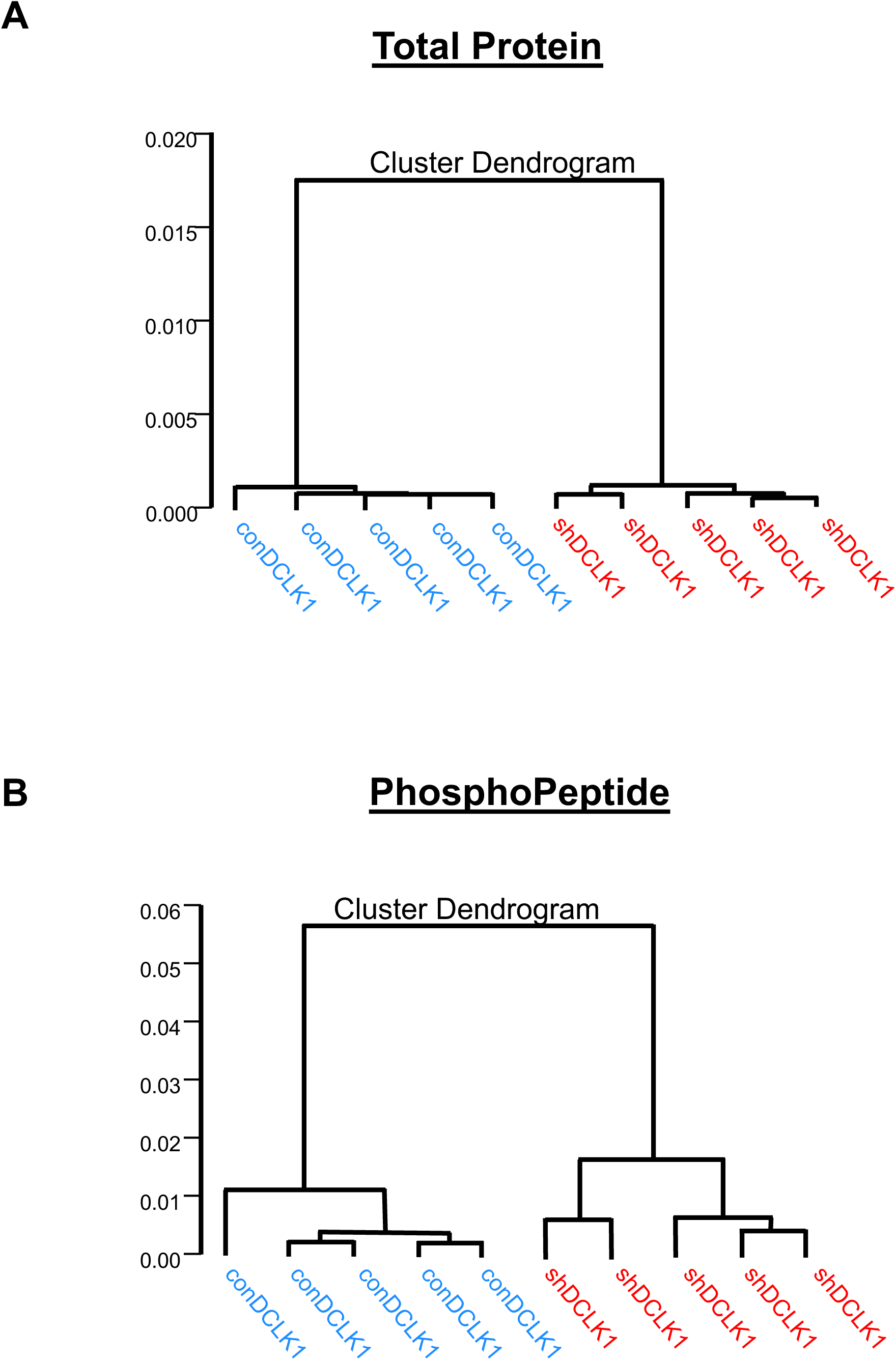
Protein expression in DCLK1 knockdown cells is distinct. Principal component analysis (PCA) was conducted on mass spectrometry data to visualize distinct clusters and dendrogram representation of both (A) total protein and (B) phosphopeptide populations. The plot exhibits clear separation between shControl (n=5) and shDCLK1 (n=5) samples, indicating distinct population patterns based on the analyzed data.

**Supplemental Figure 2.**
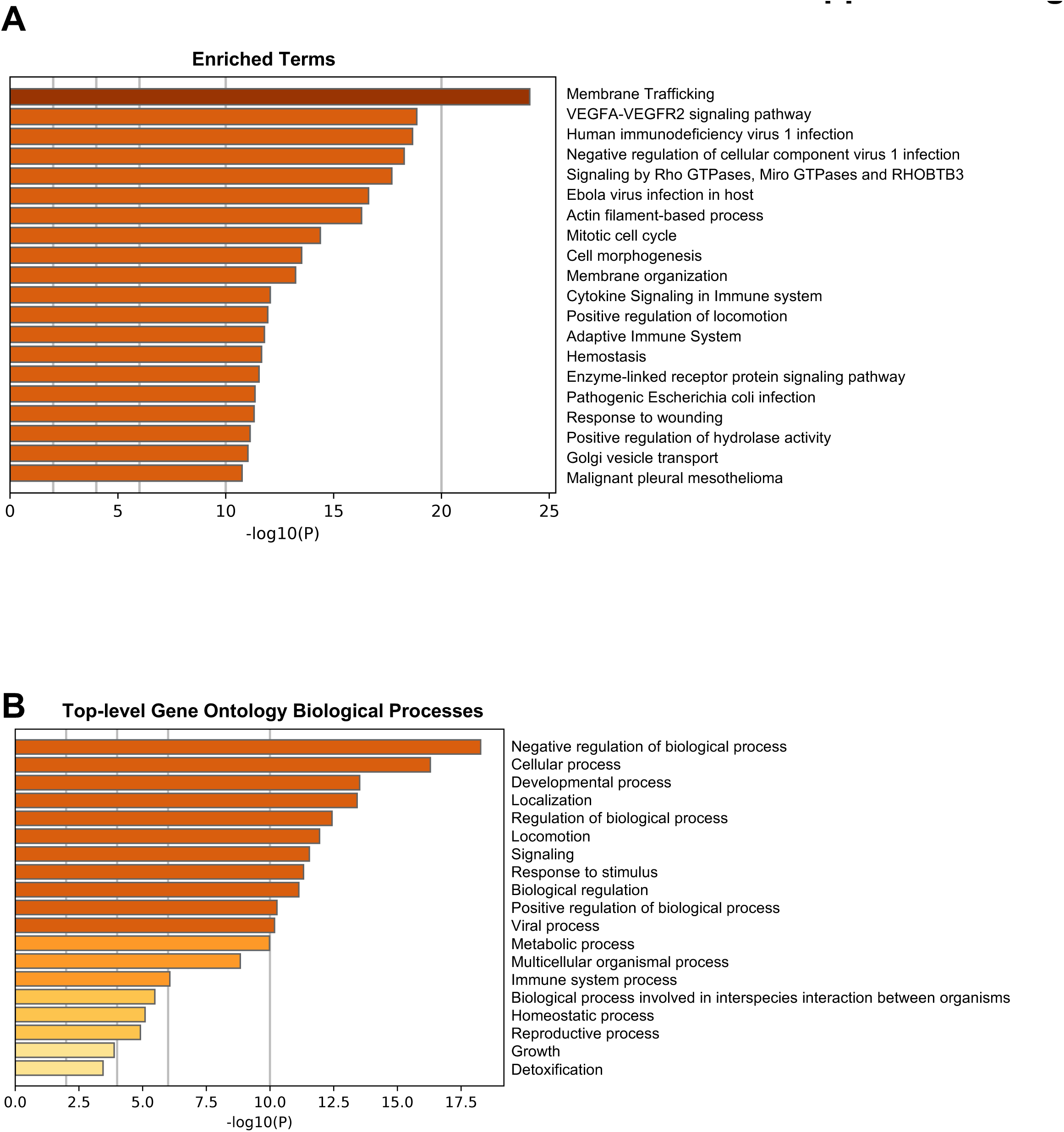
DCLK1 drives cell movement and cytoskeletal dynamics. Protein clusters displaying alterations between shControl and shDCLK1 cells were subjected to enriched ontology analysis using Metascape [18]. (A) The bar graph illustrates the top 20 clusters of Gene Ontology top-level enriched terms in biological processes, with each cluster color-coded based on its respective p-value. (B) A depiction of top-level Gene Ontology biological processes provides a comprehensive overview of the enriched terms. The analysis and images were generated using Metascape. [18].

**Supplemental Figure 3.**
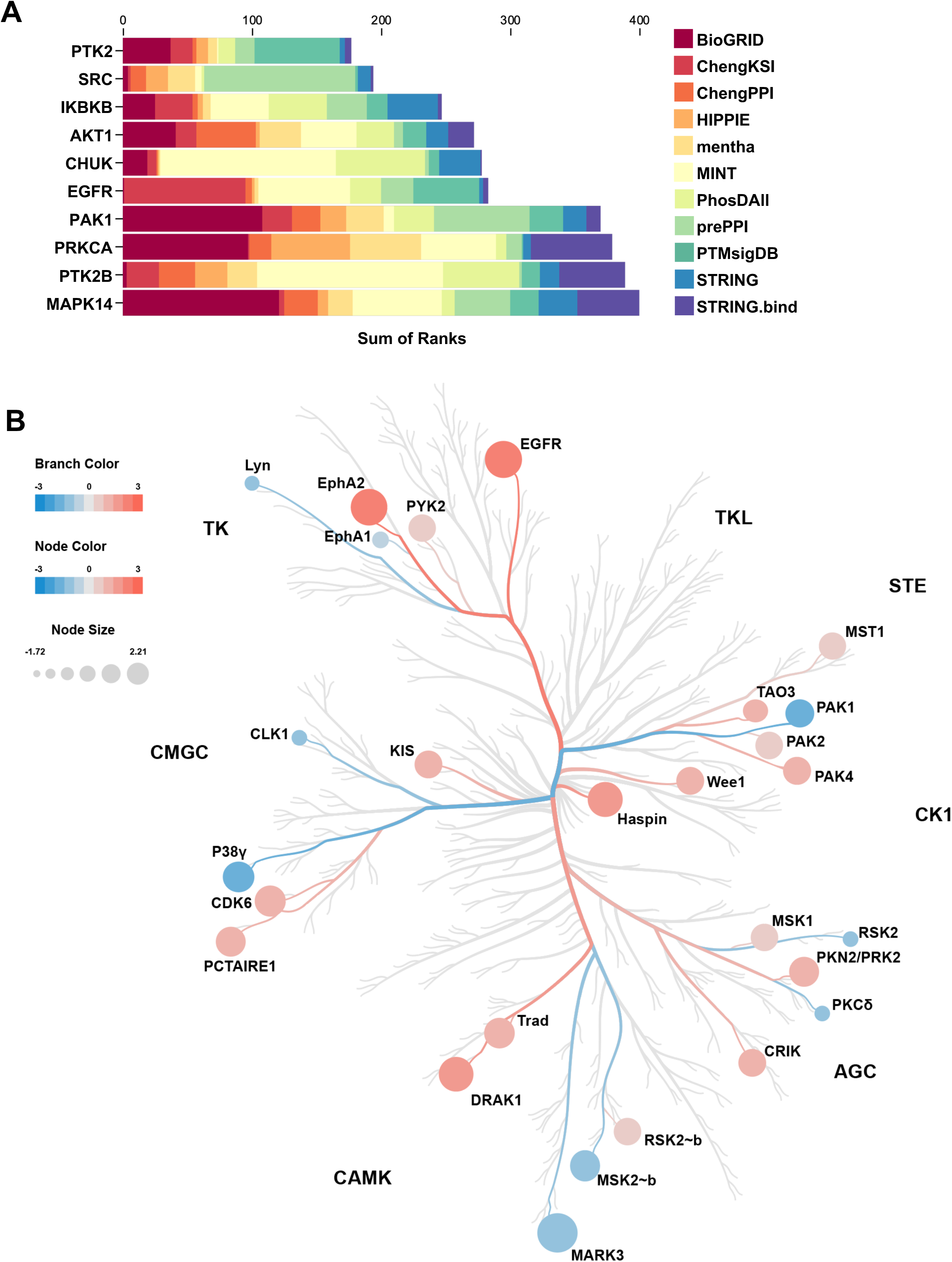
DCLK1 is associated with kinases involved in cell movement and invasion. (A) MeanRank results from KEA3 were visualized in a bar chart, emphasizing proteins enriched in shControl cells compared to shDCLK1 knockout cells. The chart presents the top ten kinases, with bars color-coded to denote the selected libraries. The integrated ranking of predicted kinases is based on MeanRank scores, and the analysis, as well as the image, was generated using KEA3. [22]. (B) A phylogenetic kinase tree is presented to compare kinase activities between the shControl and shDCLK1 groups. The tree generated using the Coral kinome tool, features nodes representing different kinases. Node color signifies protein abundance, with blue indicating increased relative abundance in the shControl group and red indicating increased relative abundance in the shDCLK1 group. The node size reflects the log2 fold change. Branches of the tree are color-coded to highlight kinase groups showing associations between the two groups. The kinome analysis and kinase tree were generated using the CORAL web app. [23].

**Supplemental Figure 4.**
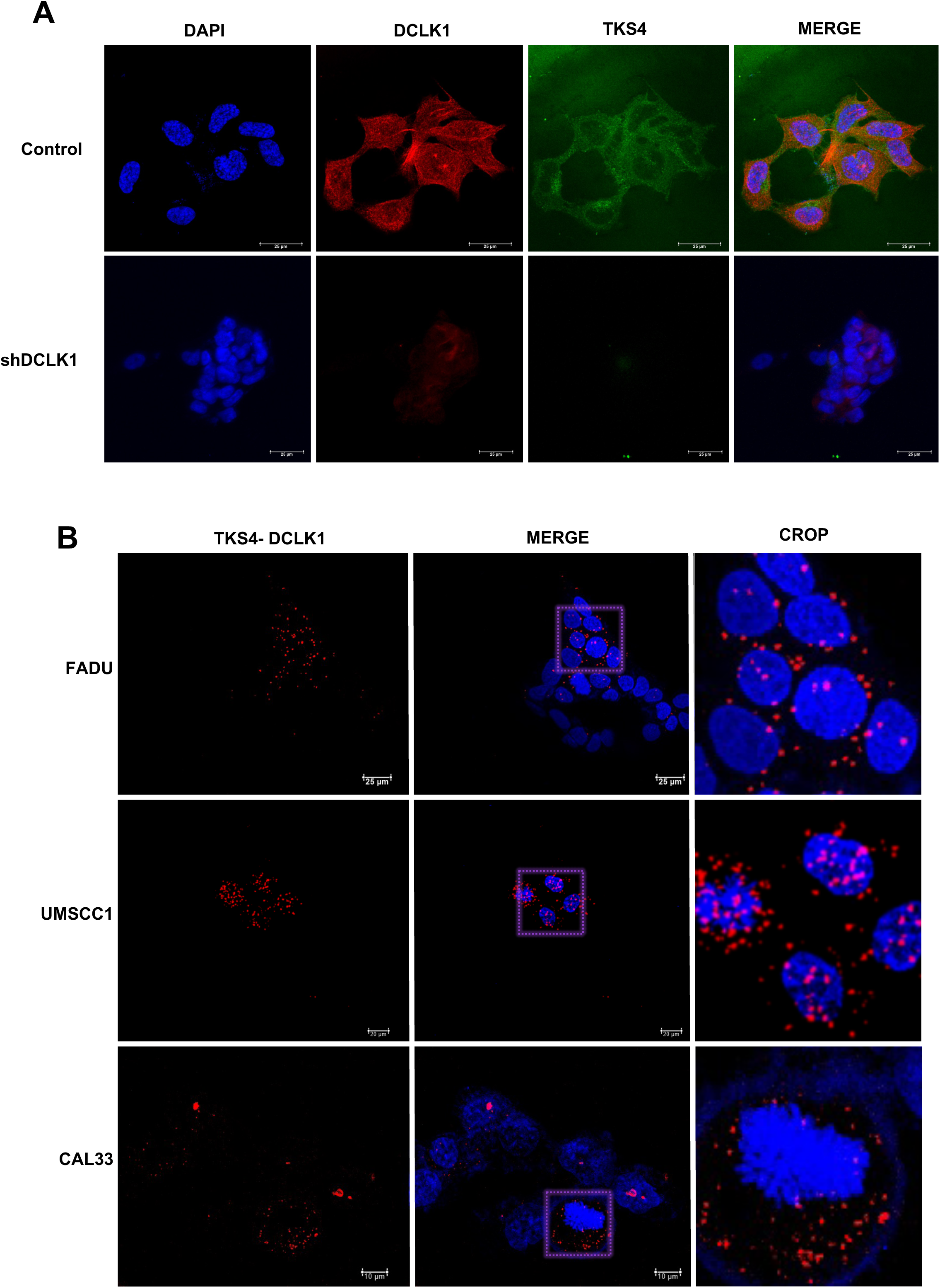
DCLK1 is associated with TKS4. (A) Confocal imaging of DCLK1 (red) with TKS4 (green) and DAPI demonstrates colocalization of DCLK1 with TKS in shControl cells, whereas such colocalization is absent in shDCLK1 FaDu cells. The images were captured with oil immersion at a 100x objective. (B) The proximity ligation assay reveals a close spatial relationship between DCLK1 and TKS4 in FaDu, UMSCC1, and CAL33 cells. Confocal imaging was performed using a 40x oil immersion objective.

**Supplemental Figure 5.**
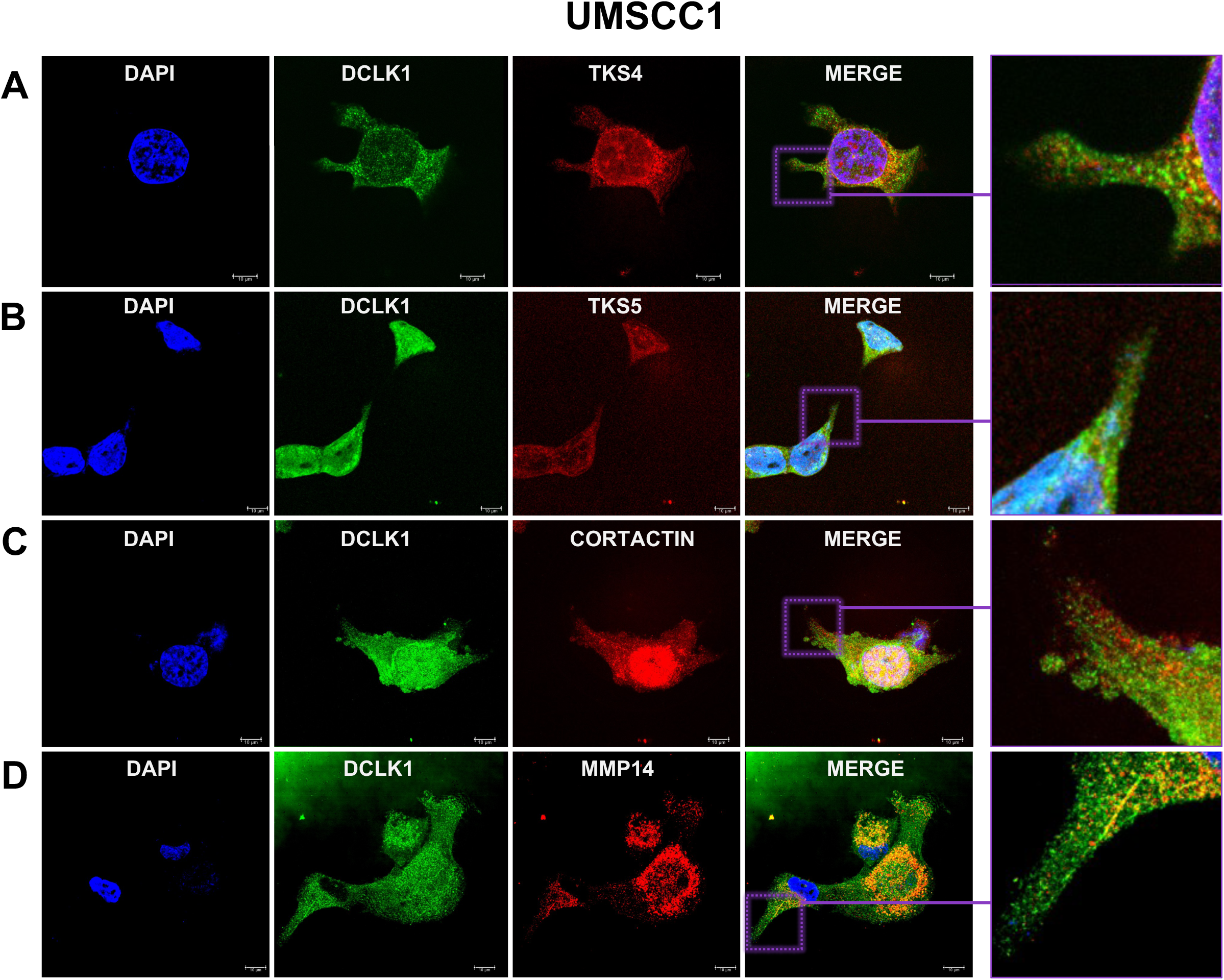
DCLK1 colocalizes with markers of invadopodia. Confocal imaging of DCLK1 (green) with invadopodia markers (red) and DAPI reveals colocalization of DCLK1 with (A) TKS4, (B) TKS5, (C) cortactin, and (D) MMP14. Oil immersion at 100x objective in UMSCC1 cells.

**Supplemental Figure 6.**
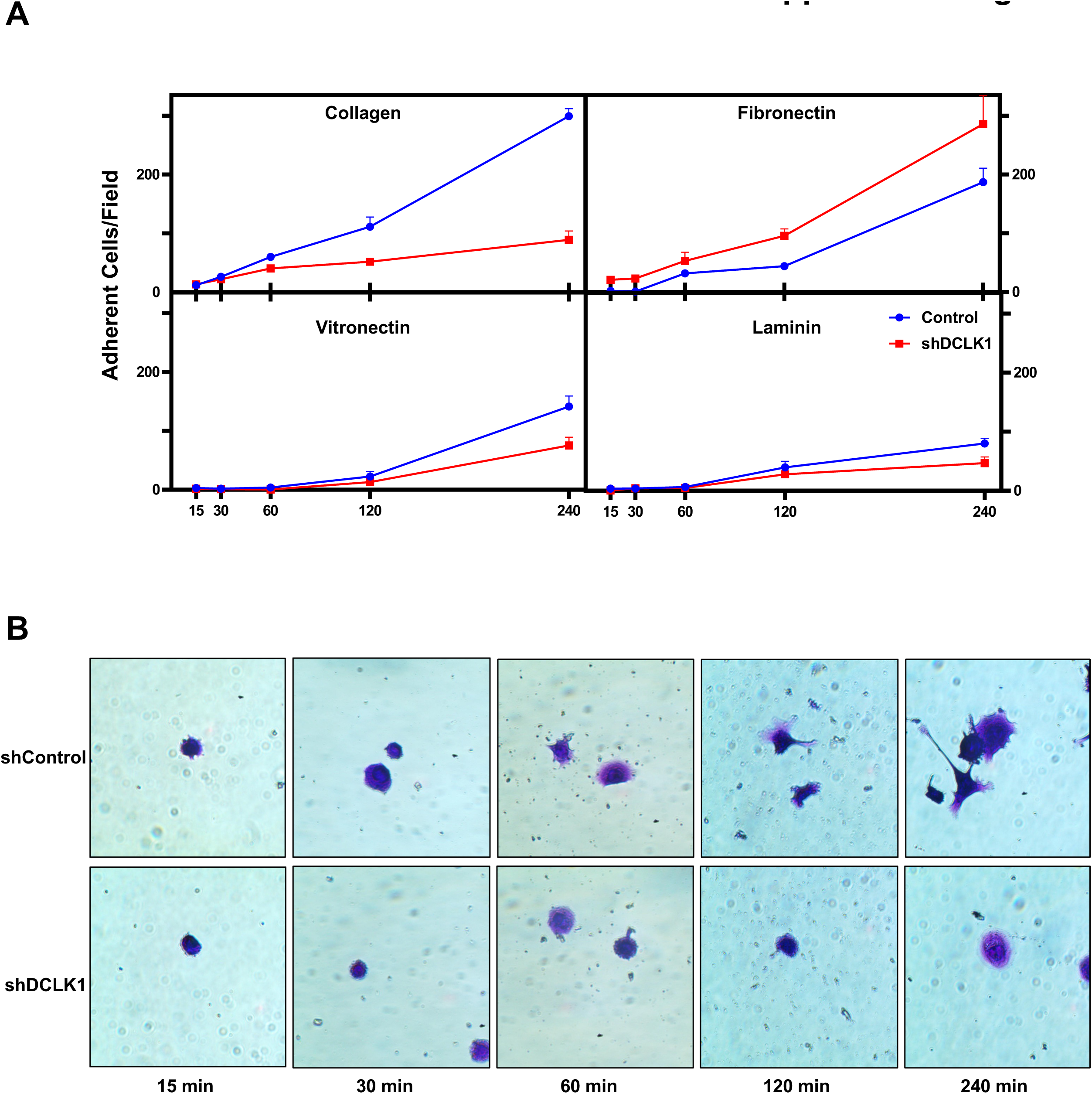
DCLK1 regulates cell adherence. (A) Adherence assay conducted on FaDu HNSCC cells, comparing shControl and shDCLK1 conditions. Knockdown of DCLK1 significantly diminishes adherence to collagen, vitronectin, and laminin over a 240-minute period, while no substantial impact on adherence to fibronectin is observed. Error bars in the graph represent ± SEM. (B) The accompanying image illustrates that DCLK1 knockdown inhibits cell spreading, as depicted in the brightfield image captured at 40x magnification using an air objective.

**Supplemental Figure 7.**
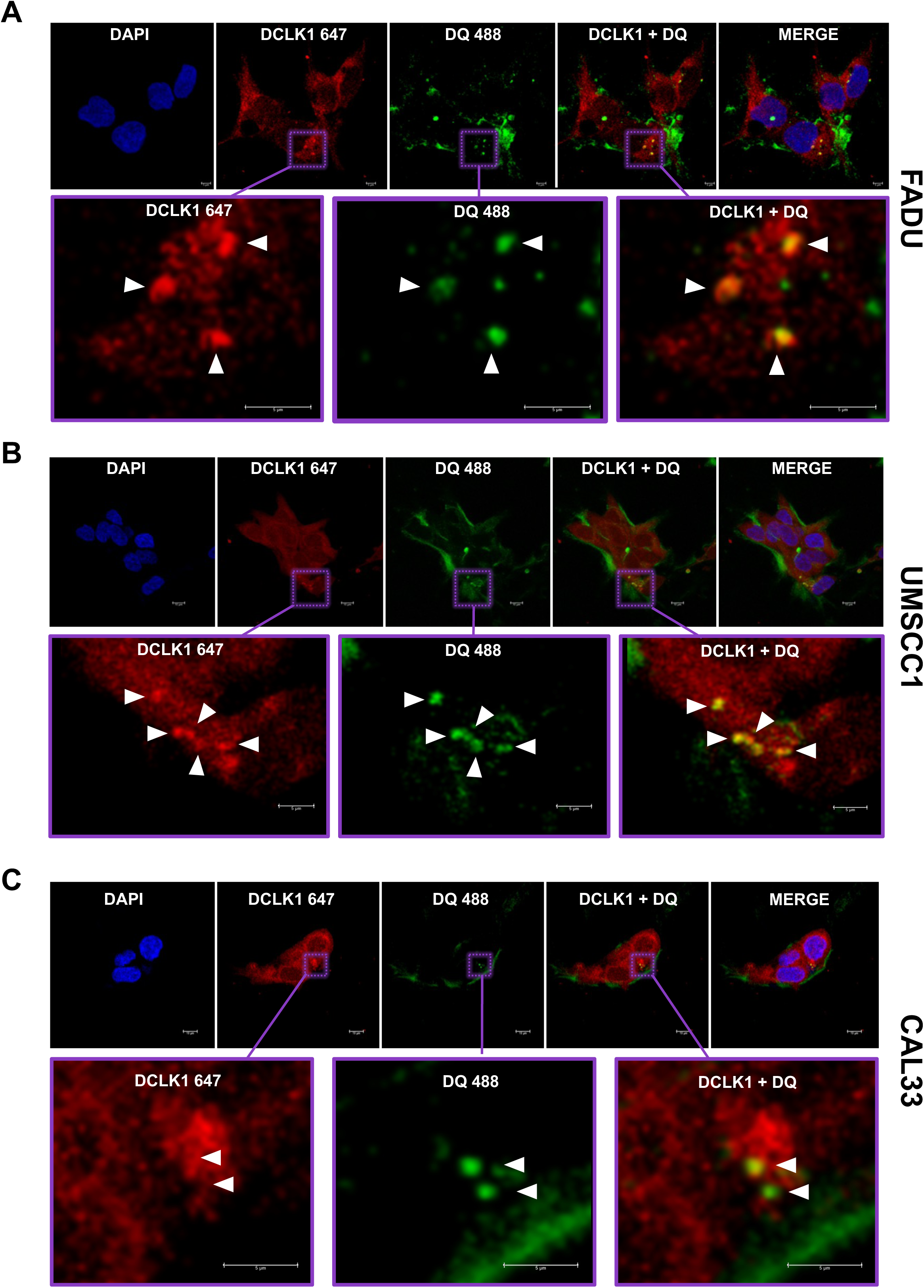
DCLK1 degrades gelatin on the ventral surface of HNSCC. Confocal imaging of (A) FaDu, (B) UMSCC1, and (C) CAL33 seeded upon DQ gelatin. Areas of DQ fluorescence degradation that overlap with increased DCLK1 expression on the ventral surface of the cells are indicated in the cropped image with white arrows. Confocal imaging performed at 60x oil immersion objective.

**Supplemental Figure 8.**
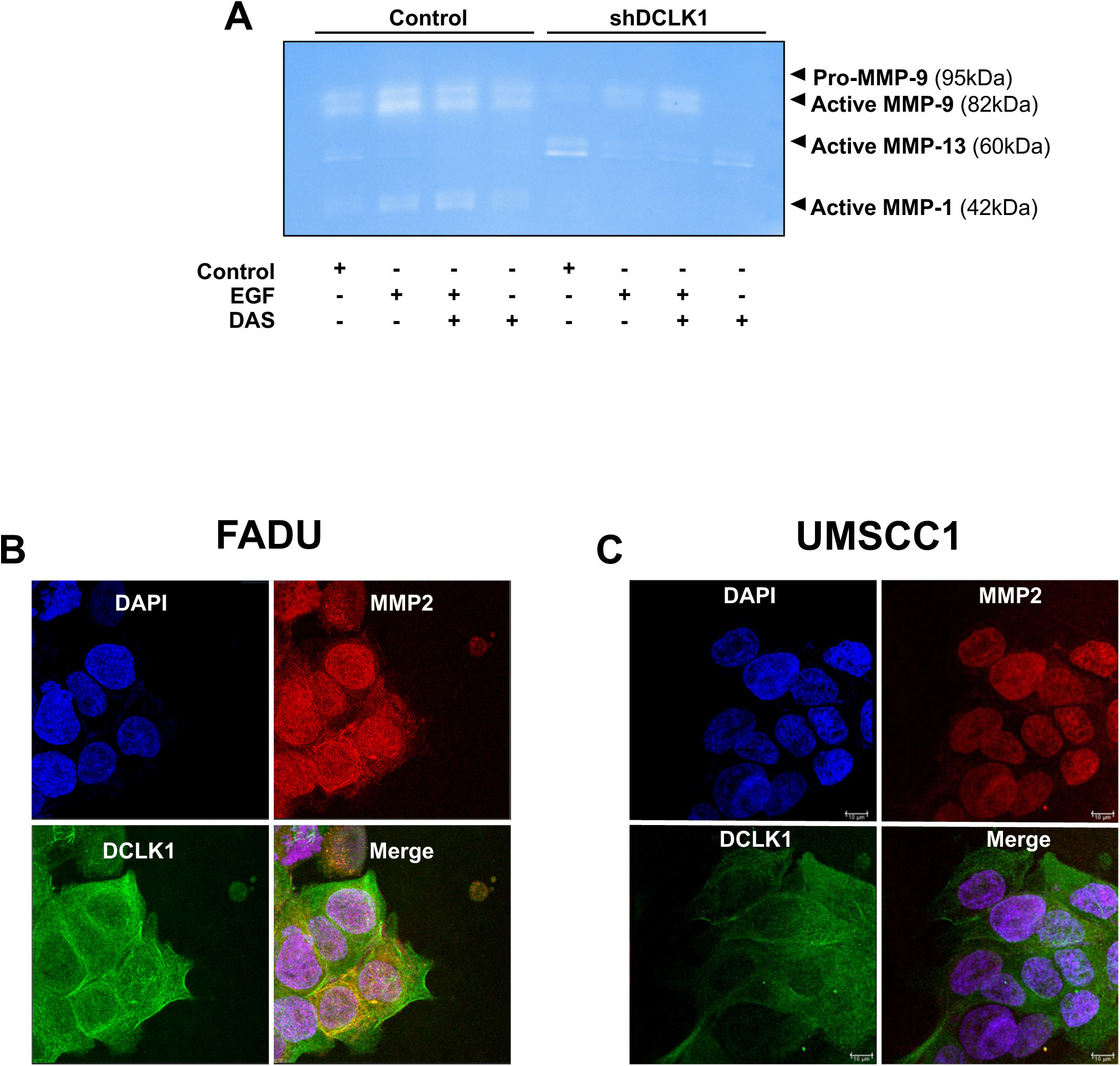
DCLK1 preferentially secretes MMP 1, 9. (A) Gel zymography analysis of concentrated supernatants from two experimental arms: shControl and shDCLK1 FaDu cells. White bands against a stained blue background indicate degradation areas, with specific MMPs migrating at known molecular weights. Each experimental arm includes variations in treatment conditions: control, EGF, EGF + dasatinib, or dasatinib. (B) FaDu and (C) UMSCC1invadopodia assay visualized via confocal microscopy. DCLK1 (green) and MMP2 (red) exhibit distinct localization patterns without colocalization or merging (yellow) in FaDu and UMSCC1 cells. Images were captured using a 40x oil immersion objective.

**Supplemental Figure 9.**
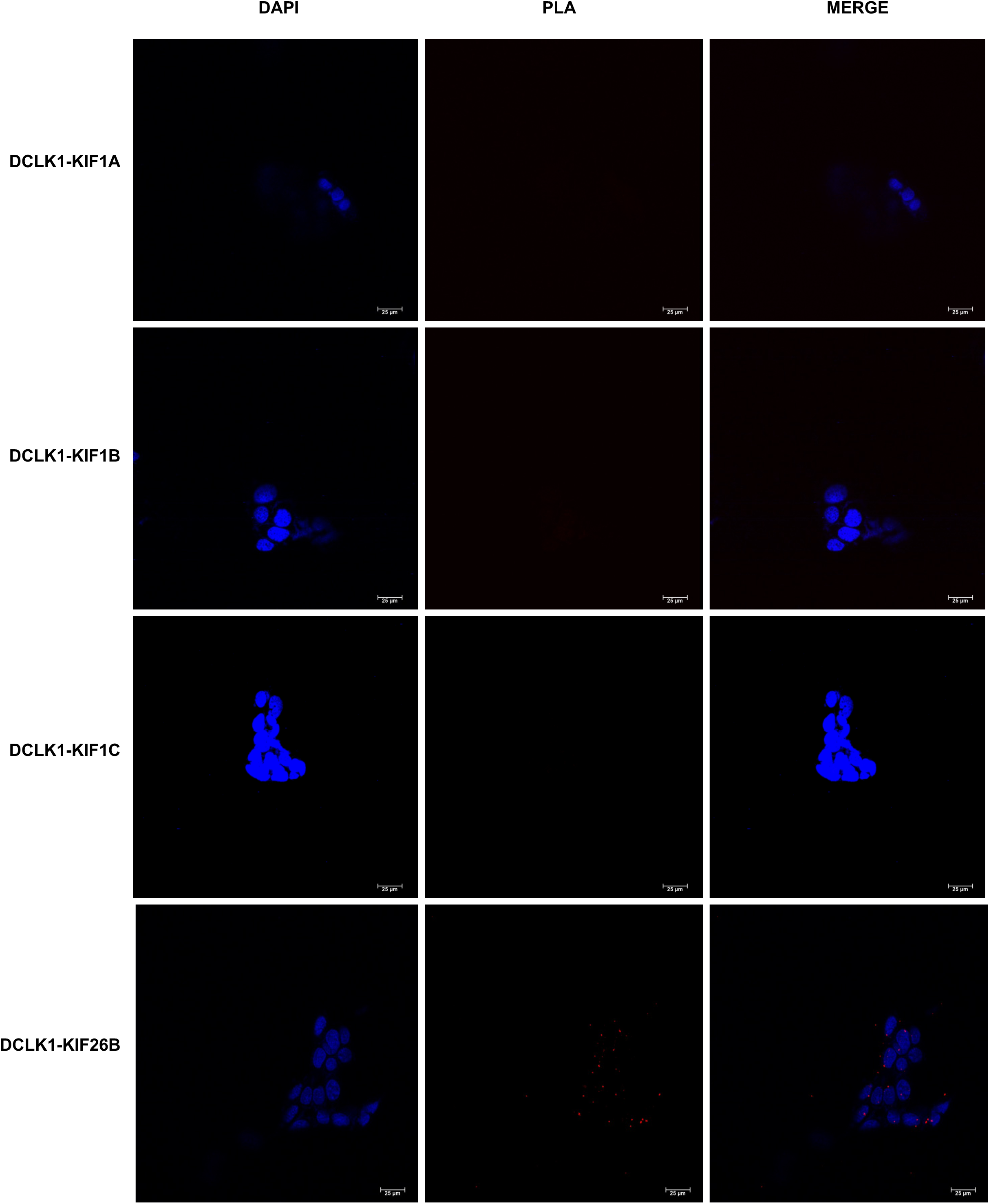
DCLK1 associates with specific Kinesin 3 family members. PLA screening of the kinesin 3 family members with DCLK1 in FaDu cells reveals minimal to no association between DCLK1 and KIF1A, KIF1B, KIF1C, or KIF26B. Oil immersion at 40x objective. DAPI nuclear stain and red puncta indicate PLA interactions.

**Supplemental Figure 10.**
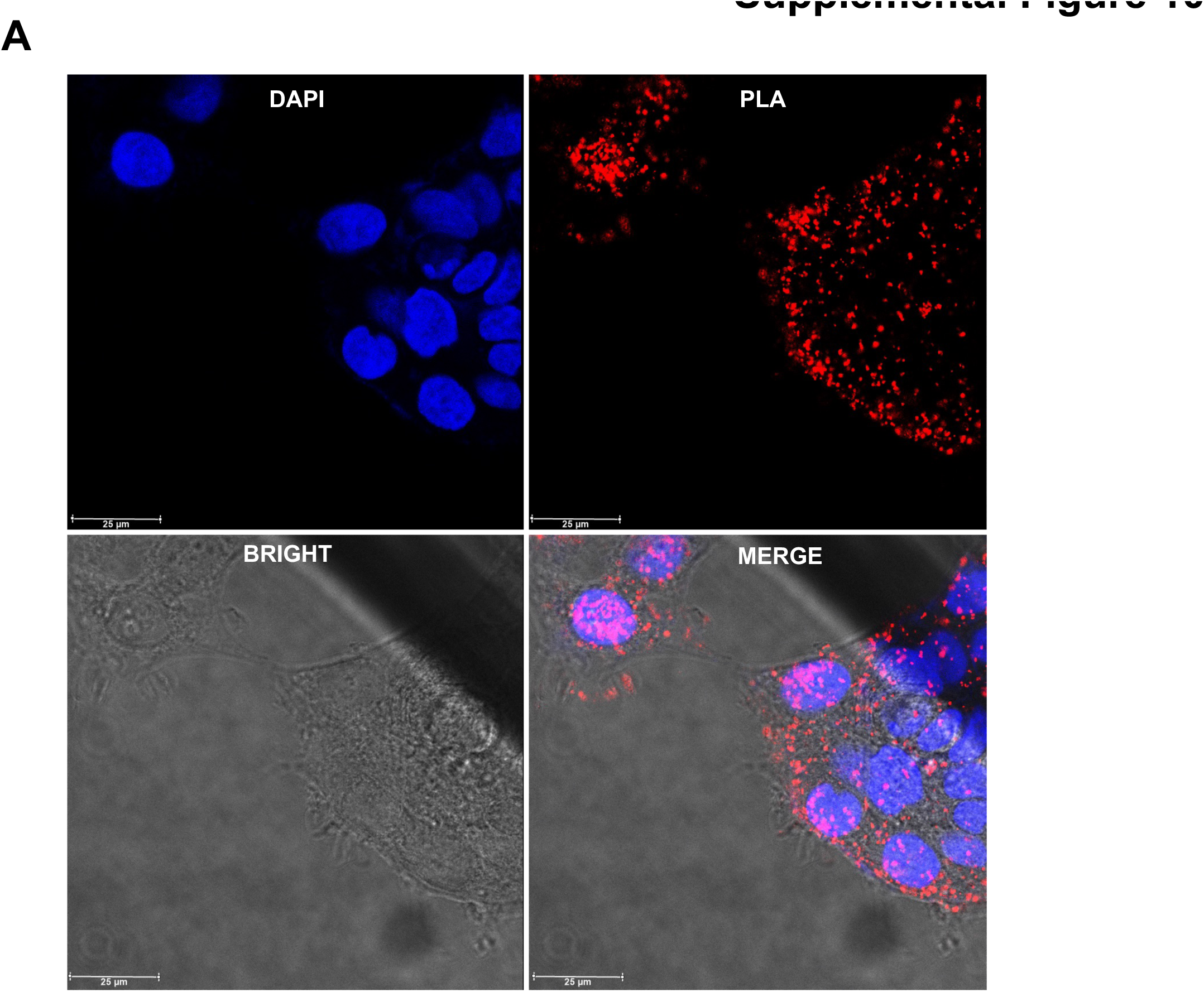
DCLK1 strongly associates with KIF16B. PLA visualization of DCLK1 and KIF16B in FaDu cells, presented without cropping, as depicted in Figure 5E. The imaging was performed using a 40x oil immersion objective.

**Supplemental Figure 11.**
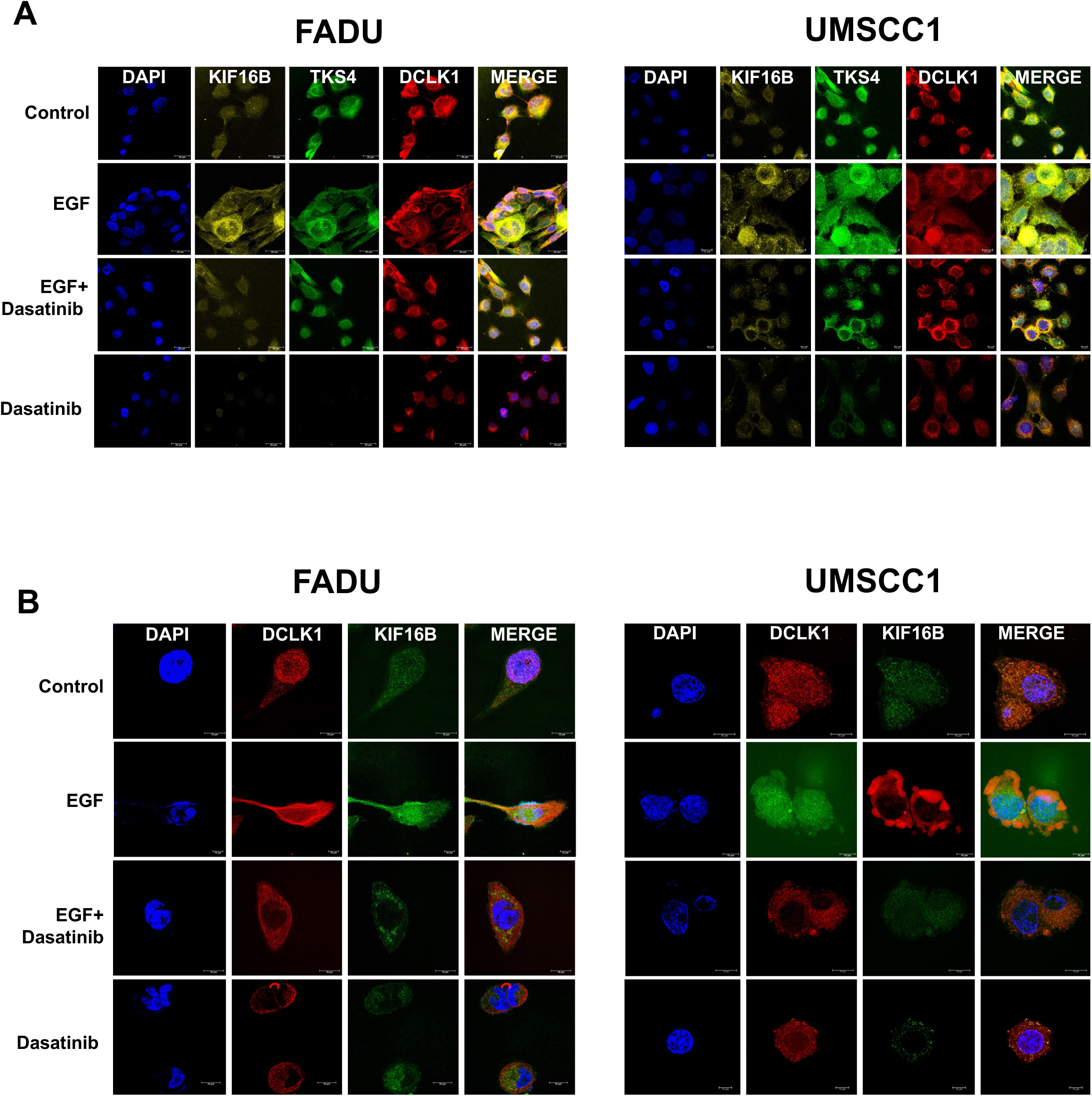
DCLK1 colocalization to KIF16B is modulated by EGF/Dasatinib. Confocal microscopy images depict DCLK1 (red), KIF16B (yellow), and TKS4 (green) in FaDu and UMSCC1 cells. Notably, enhanced colocalization of DCLK1 with KIF16B and TKS4 is observed following EGF (10 ng/ml) stimulation, while a reduction is evident with dasatinib (100nM) treatment. The accompanying bar graph highlights significant differences in colocalization (merge) during stimulation or inhibition. Images were captured using oil immersion at (A) 40x and (B) 100x objectives.

**Supplemental Figure 12.**
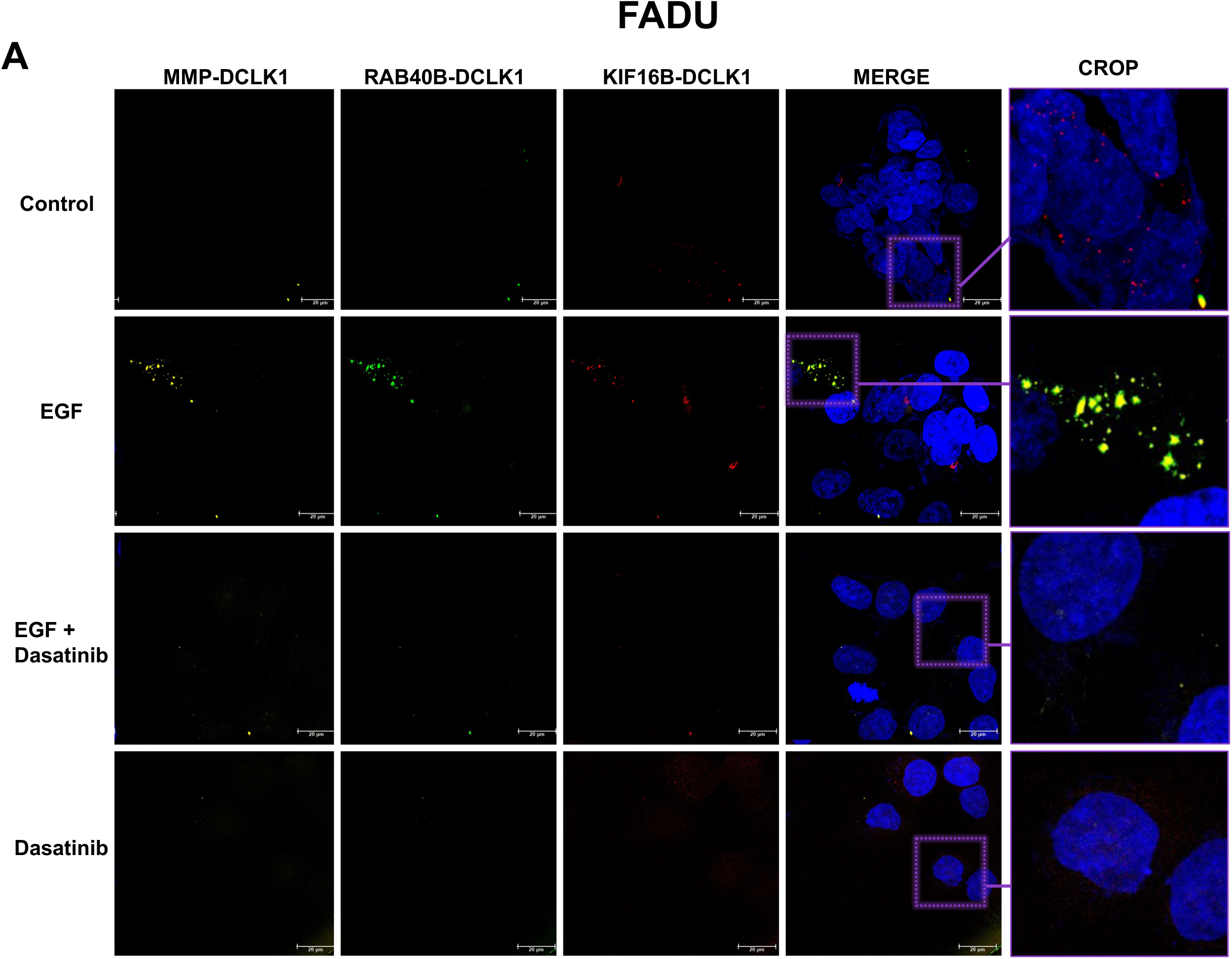
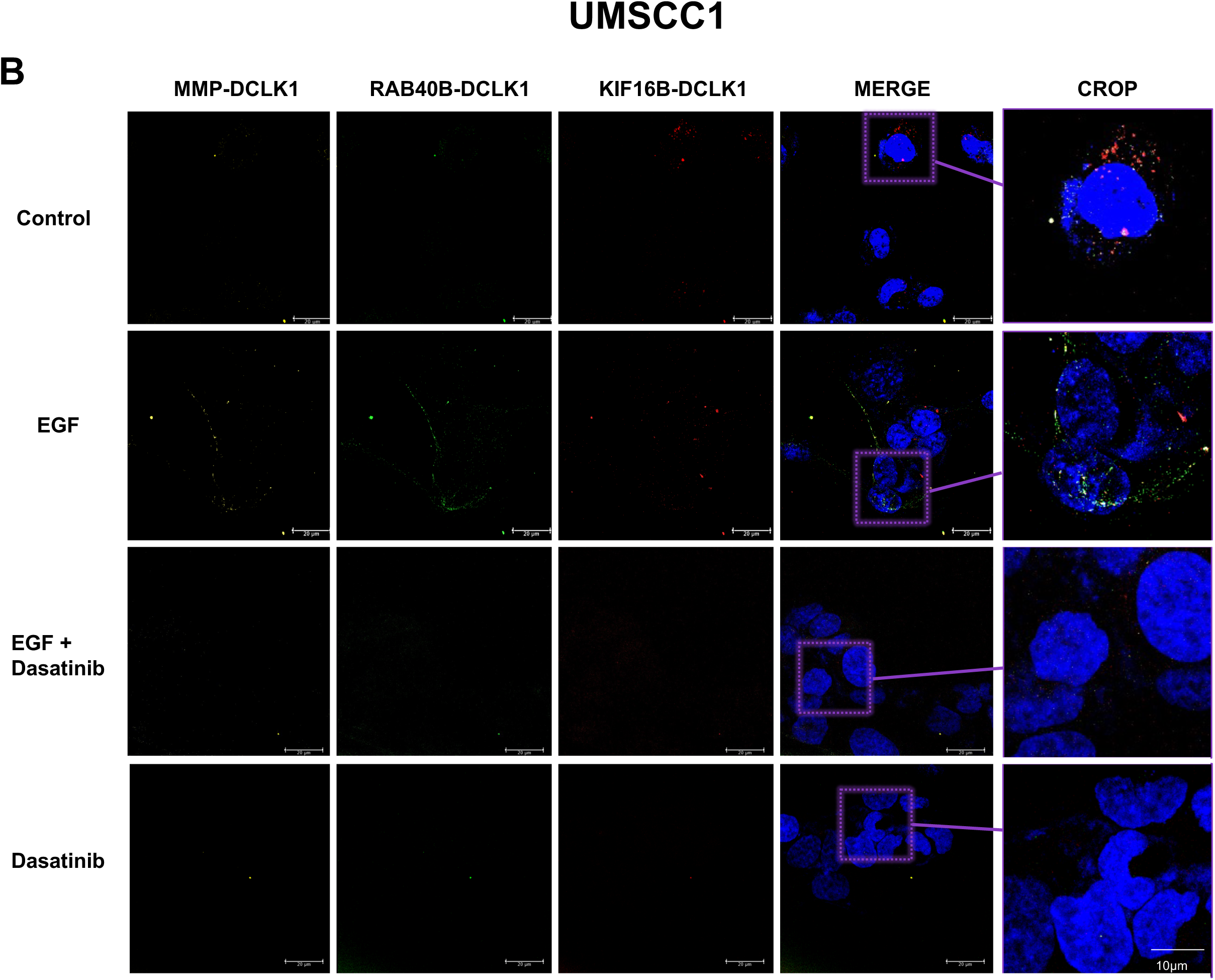
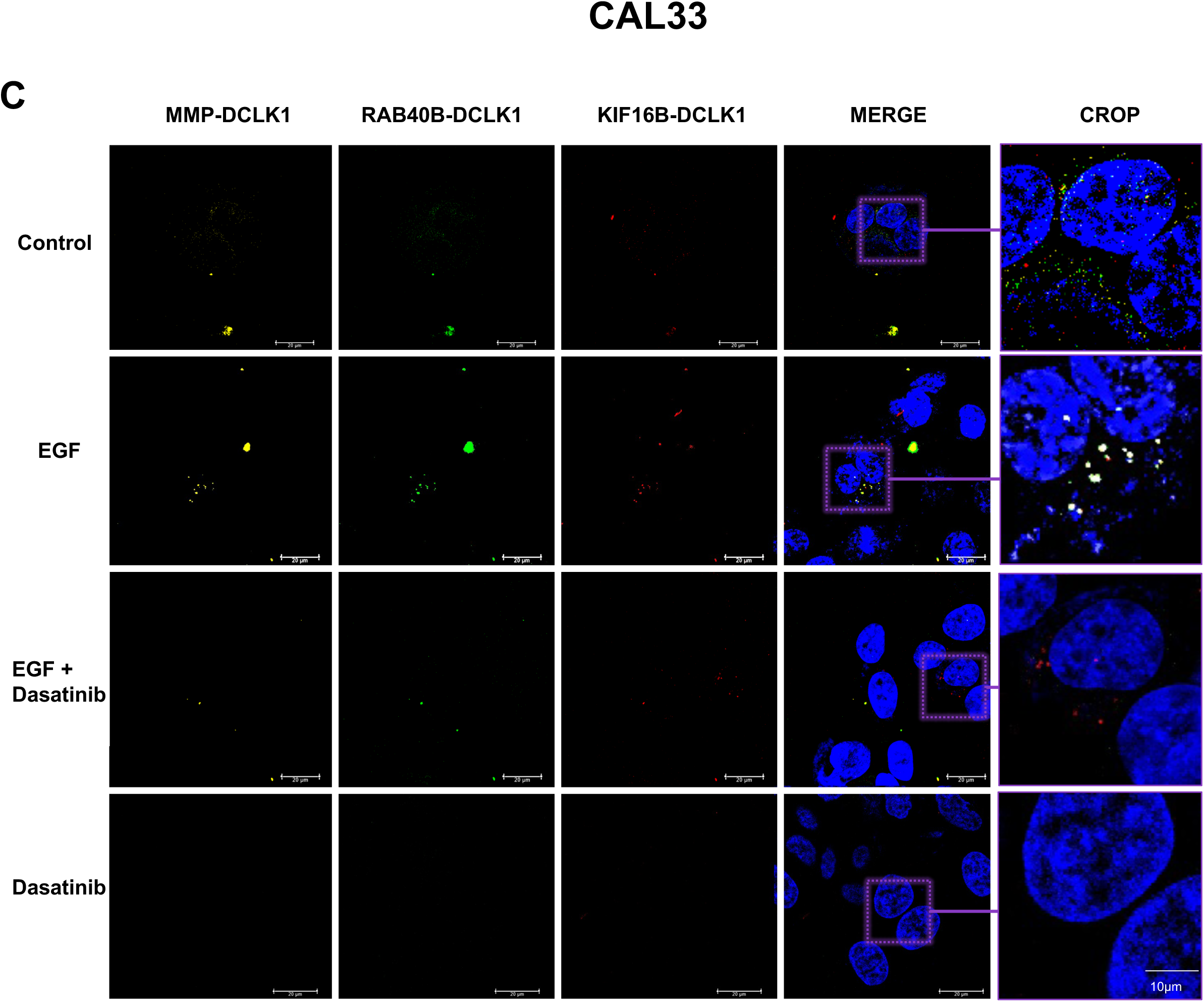
DCLK1, MMP9, RAB40B, and KIF16B form a complex. PLA was conducted in (A) FaDu, (B) UMSCC1, and (C) CAL33 to visualize the interactions between DCLK1 and MMP9, RAB40B, and KIF16B family members. Nuclear staining was performed using DAPI, and positive interactions were indicated by red puncta in the proximity assay PLA. Images were captured with oil immersion at a 40x objective.

**Supplemental Figure 13.**
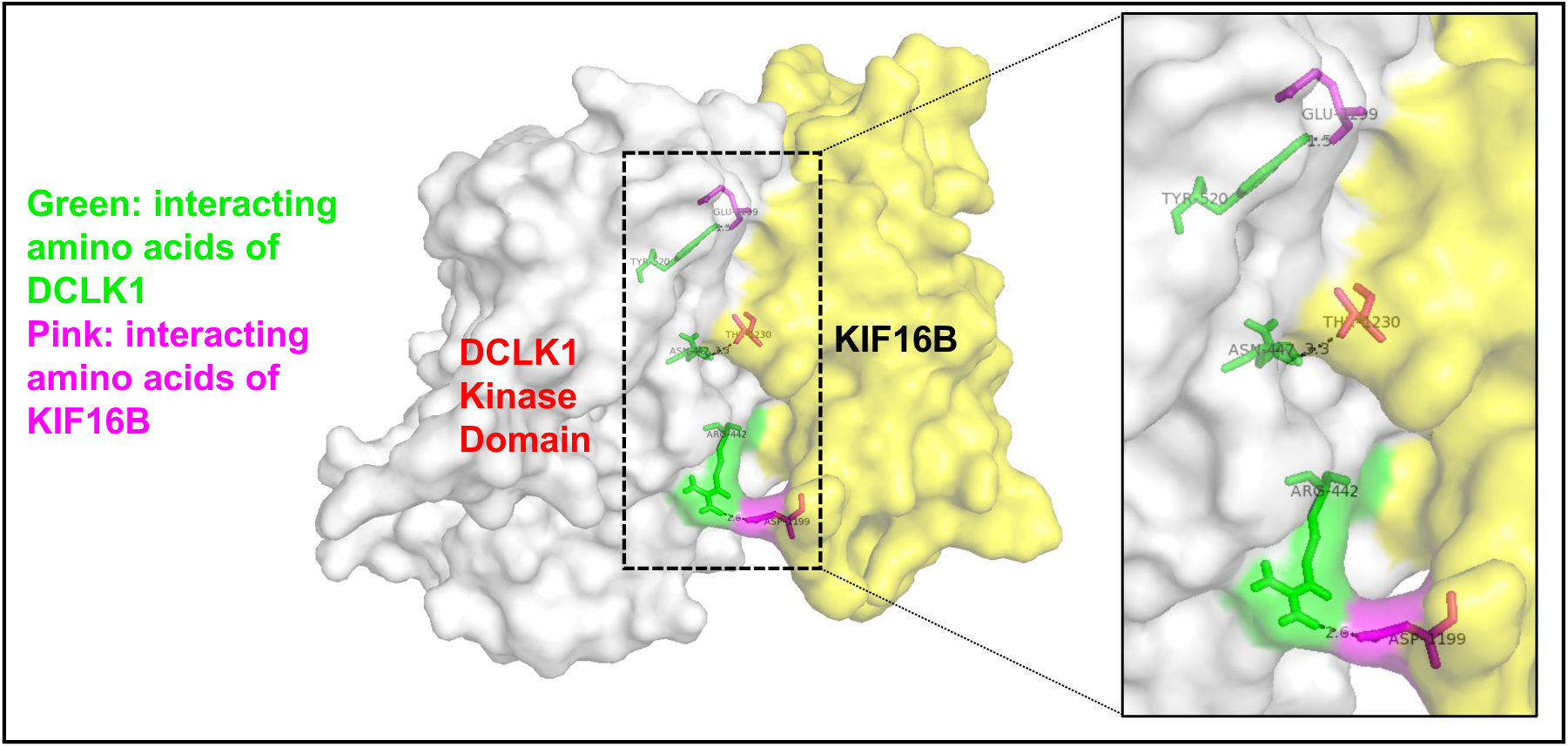
Predictive analysis demonstrate DCLK1-KIF16B binding. Computational analysis of the binding interaction between the DCLK1 kinase domain and KIF16B motor domain revealed a specific interaction site at arginine 442 in DCLK1 and aspartic acid 1199 in KIF16B, with a distance of 2.6 angstroms.

